# Comprehensive characterization of the complex BAHD acyltransferase family from 218 land plants species: phylogenomic analysis and identification of specificity determinant positions

**DOI:** 10.1101/2023.03.15.532435

**Authors:** Fernando Villarreal, Agustin Amalfitano, Hugo Marcelo Atencio, Nicolas Stocchi, Arjen ten Have

## Abstract

Chemodiversity is a fundamental trait acquired by plants during their land’s colonization. This resulted from an evolutionary process leading to the increase in the number of homologues from a distinct set of protein superfamilies, many of them associated to the specialized metabolism, which allowed the expansion of the chemical space to cope with several environmental cues. BAHD acyltransferases are among these important superfamilies, catalyzing a reaction leading to the acylation of acceptor metabolites with Coenzyme A-activated donors. BAHD acyltransferases can use a wide variety of substrates and they often times display substrate permissiveness towards a wide variety of substrates. Together, these factors complicates the reliable identification and functional annotation of BAHD homologues, also due to the (relatively) limited amount of biochemical data on BAHD acyltransferases. In this work, we take a phylogenomics and computational approach to study the BAHD superfamily in land plants. Using a clustered training set with 27 proteomes, followed by the classification of additional 191 proteomes, we obtained a final BAHDome with 15607 homologues. The training set was clustered in 16 groups, that, together with the identification of cluster of orthologues from the complete BAHDome, were partially assigned functionally to different families guided by the characterized activities (in terms of metabolites used) present in each group. However, the function assignation was not direct in several cases, due to large sequence number, taxonomical distribution on the group, and high sequence variability intra-group. Finally, we used the different families (and functional subfamilies) identified to detect specificity determining positions (SDPs), that may account to explain the functional diversity by finding key positions in, for example, substrate interaction. However, only a handful of the SDPs identified in this work are linked to the substrate binding pocket. Together, these results allow to partially annotate functional BAHD families, which have evolved in a complex pattern of taxonomical and functional signals to allow interaction with multiple substrates, more likely associated to protein dynamics rather than direct substrate interaction.

## INTRODUCTION

Specialized metabolism is a term for the pathways and small metabolites that are not absolutely required for the survival of the organism. Only a few examples of specialized metabolites are pigments as anthocyanins and carotenoids, volatiles such as terpenes, alkaloids and glucosinolates. Many specialized metabolites are precursors for polymers as lignin and suberin, other components contribute to smell and taste and/or are involved in interaction with the environment and are important in stress mechanisms [1], [2]. Specialized metabolism and its metabolites form what is referred to as phytochemistry and which is renowned for its complexity and potential for medical applications in the form of nutraceutical compounds or medicine [3], [4].

How metabolic pathways evolved is largely an open question. However, analyses of specialized metabolic subnetworks suggest that functional redundancy and diversification has been playing an important role [5][6]. Typically, the different networks comprising the specialized metabolism have most likely evolved from a simple set of (rather) linear reactions, similar to what has been shown for microbial metabolic networks [7]. Gene duplication generate a functional redundancy that allows for functional diversification, which is facilitated by the fact that specialized metabolism in general is under a lower functional constraint than central metabolism. This facilitated by the high robustness shown by protein folding in specialized metabolism superfamilies, As such, this results It is also clear that functional redundancy contributes to the robustness of the network [5]. This is accompanied by the notion that mechanisms of enzyme promiscuity (mainly substrate permissiveness [8]) are essential for evolution of new functions (e.g., new metabolic pathways) [9].

The BAHD acyltransferase superfamily (named according to the first letter of each of the first four biochemically characterized enzymes of this family, to wit: **B**EAT: benzylalcohol O-acetyltransferase; **A**HCT: anthocyanin O-hydroxycinnamoyltransferase; **H**CBT: anthranilate N-hydroxycinnamoyl/benzoyltransferase; and **D**AT: deacetylvindoline 4-O-acetyltransferase) contributes with key components in different pathways of the specialized metabolism [10]. BAHD enzymes are typically cytoplasmic, soluble proteins and ∼50 kDa, catalyze the transfer of a acyl moiety activated by Coenzyme A (CoA), referred to as donor, to an acceptor metabolite via the creation of either ester or amide bonding, although this mechanism may be different for some of the BAHD enzymes [11]. Classically, BAHD enzymes are described to possess a motif HxxxD, buried in the protein and key on the enzyme’s catalytic process, and a second motif DFGWG, which is not located in the proximity of the binding pocket [12].

With the advent of almost every genome sequence, protein superfamilies can also be studied by computational biology. Protein sequence analysis, but also protein structure modeling [13] have developed a number of reliable methods for structure-function prediction, which in turn depend on the construction of high quality multiple sequence alignments, together with phylogenetic reconstruction for protein superfamilies. These methods essentially concerns a superfamily study, that typically consists of several subfamilies, sequences from somewhat distant homologues that need to be classified. For this, we will use the recently developed HMMERCTTER [14] since it has been shown to outperform the Panther phylogenomics suite on a number of protein superfamilies.

One important type of characteristic that is used to explain functional differences among families are specificity determining positions (SDPs), that can be detected by various approaches. The basics of most approaches is best known as evolutionary tracing [15], [16], which uses a phylogenetic hierarchy and corresponding multiple sequence alignment (MSA) to identify positions that show conservation at a certain hierarchic level. Other methods such as SDPfox [17] use mutual information (MI) in between MSA columns and clustering in order to identify Clustering Determining Positions (CDPs). Since selection of substitutions can occur as a result of drift rather than selection, these are not necessarily SDPs. Other MI methods such as Mistic [18] compare the columns of an MSA. High MI between two MSA columns suggests the corresponding positions somehow interacted during evolution. We combine SDPfox with Mistic below the premise that CDPs that show high MI are unlikely to have evolved by drift. We apply this using a network approach and a network connectivity score in order to select the most probable SDPs. We have ap pC lied this approach to the identification of SDPs in other families [19][20].

To the best of our knowledge, there are no studies at the phylogenomic level across land plants available. As such, we aim to understand phylogenetic history of plant BAHD utilizing a wide dataset of 218 complete proteomes, and use it to clusterize the superfamily into families showing potential functional diversification. Guided by characterized enzymatic activities, we also will attempt to find links between families and substrates selectivity, and finally identify key residues that may be associated to functional diversification, hence SDPs, which may be associated to binding pocket to allow substrate selectivity.

## RESULTS

### Sequence mining identifies 1761 BAHD homologue sequences in 27 land plants

We performed a sensitive data mining using a HMMER profile for BAHD family from Pfam (PF02458). The profile was used to retrieve homologues from 27 land plants proteomes (**Figure 1**), including two bryophytes (*P. patens* and *M. polymorpha*), one lycophyte (*S. moellendorffii*); two ferns (*A. filliculoides* and *S. cucullata*); three gymnosperms (*G. biloba* and two Pinales, *T. baccata* and *P. abies*); and 19 angiosperms. The latter includes five monocot species (one Araceae, *S. polyrhiza*, and four Poaceae: *Z. mays, S. bicolor, B. distachyon* and *O. sativa*), and 13 eudicot species: *A. hypochondriacus* for Caryophyllales, three Asterids (*H. annus, S. tuberosum* and *S. lycopersicum*), and nine Rosids (four fabids *M. truncatula, P. persica, M. esculenta, P. trichocarpa*, four malvids *B. rapa, A. thaliana, T. cacao, C. sinensi*, and incert classified *V. vinifera*). We also included *A. trichopoda*, which is a sister species to all angiosperms. In addition, we retrieved all Viridiplantae entries from UniProtKB/SwissProt (SP) database with highest curation levels (1 & 2), and we treated it as an independent proteome for homologue search. In addition, we included to these SP database 39 entries from SP lower curation levels, previously described in the literature as functional BAHD [21].

**Figure 1:**
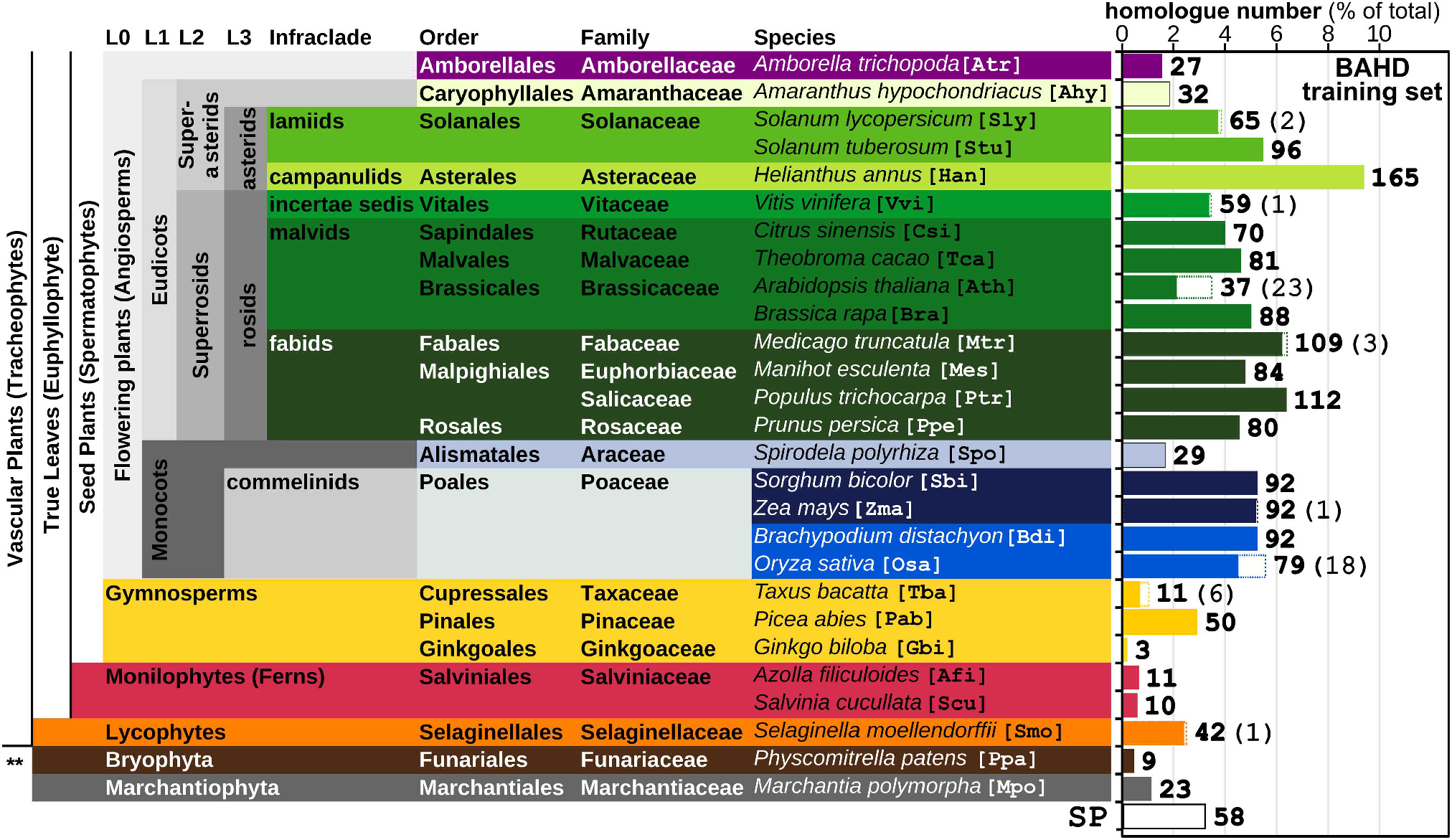
Schematic view of taxonomic contribution of *BAHD homologue sequences*. Taxonomic arrangement of the 27 land plants selected for the study. Three letter code for each species is denoted in square brackets. L0-L3 denote different clade hierarchical levels for Angiosperms. Plot to the right shows percentage of BAHD homologues per species identified in the final dataset. White boxes indicate UniProtKB/SwissProt (SP) entries corresponding to the given species, the SP bar below corresponds to SP entries from other species. Numbers on the plot indicate actual number of BAHD homologues per species (the amount of SP entries in brackets). **, Bryophyte.

The sensitive sequence mining yielded a total of 2891 putative BAHD homologue sequences, including 114 SP entries, and ranging from 17 sequences in *A. filliculoides* to 216 in *H. annus* (Supplementary Dataset S1). Next, potential non functional homologues were excluded with Seqrutinator, resulting in a total of 1748 homologues (removing 1143 sequences, 7 of them from SP). Note that 84.2% of the sequences removed correspond to partial sequences. Indeed, modules targeting partial sequences removed elements with an average length of 218.2 residues (standard deviation 106.2), compared to the average length of 453.5 (s.d. 72.9) observed for the accepted sequences. Next, a dedicated analysis of the scrutinized sequences allowed us to recover a total of 86 homologues, including six SP sequences. In a final manual step, we removed sequences lacking the BAHD motif HxxxD as well as duplicated sequences (100% identity). In the 55 cases we identified an homologue from a complete proteome with a SP counterpart, we conserved the latter, as it will be instrumental for further functional annotation. This was the case of, for example, 23 *Arabidopsis* entries also identified in SP. As a final result, we obtained a dataset of 1761 BAHD homologues (113 SP entries, 58 of them from species not present in our dataset) from 27 complete proteomes (**Figure 1**).

In general, we observe an increased BAHD homologue number in superior land plants. *H. annus* is the species with the highest number of BAHD homologues (165), followed by *P. trichocarpa* and *M. truncatula. G. biloba, P. patens* and fern *S. cucullata* presented the lowest number of homologues. A total of 32 sequences belong to Bryophytes, 42 to Lycophyte (one of them is a SP entry), 21 to ferns and 70 sequences are from Gymnosperm (6 from SP). Focusing on angiosperms, we identified 27 homologues in the basal *A. trichopoda*, 403 homologues in monocots and 1107 in eudicots.

### The land plant BAHD superfamily clusters into 16 major families organized in 7 ancestral clades

The identified sequences were used to generate a maximum likelihood phylogenetic tree, that was subsequently subjected to clustering using HMMERCTTER (**Figure 2**). The tree shows a high degree of statistiacal support for most branches, as shown by the transfer bootstrap expectation (TBE) values (**Fig 2A**). HMMERCTTER clustering results in 16 clusters with 100% precision and recall self detection (100P&R-SD). This basically means that each cluster shows higher internal rather than external similarity. This similarity based clustering is conceptually the result of taxonomic effects or functional diversification. Note that groups are termed G1 to G16, being the largest to smallest groups in terms of number of homologues.

**Figure 2:**
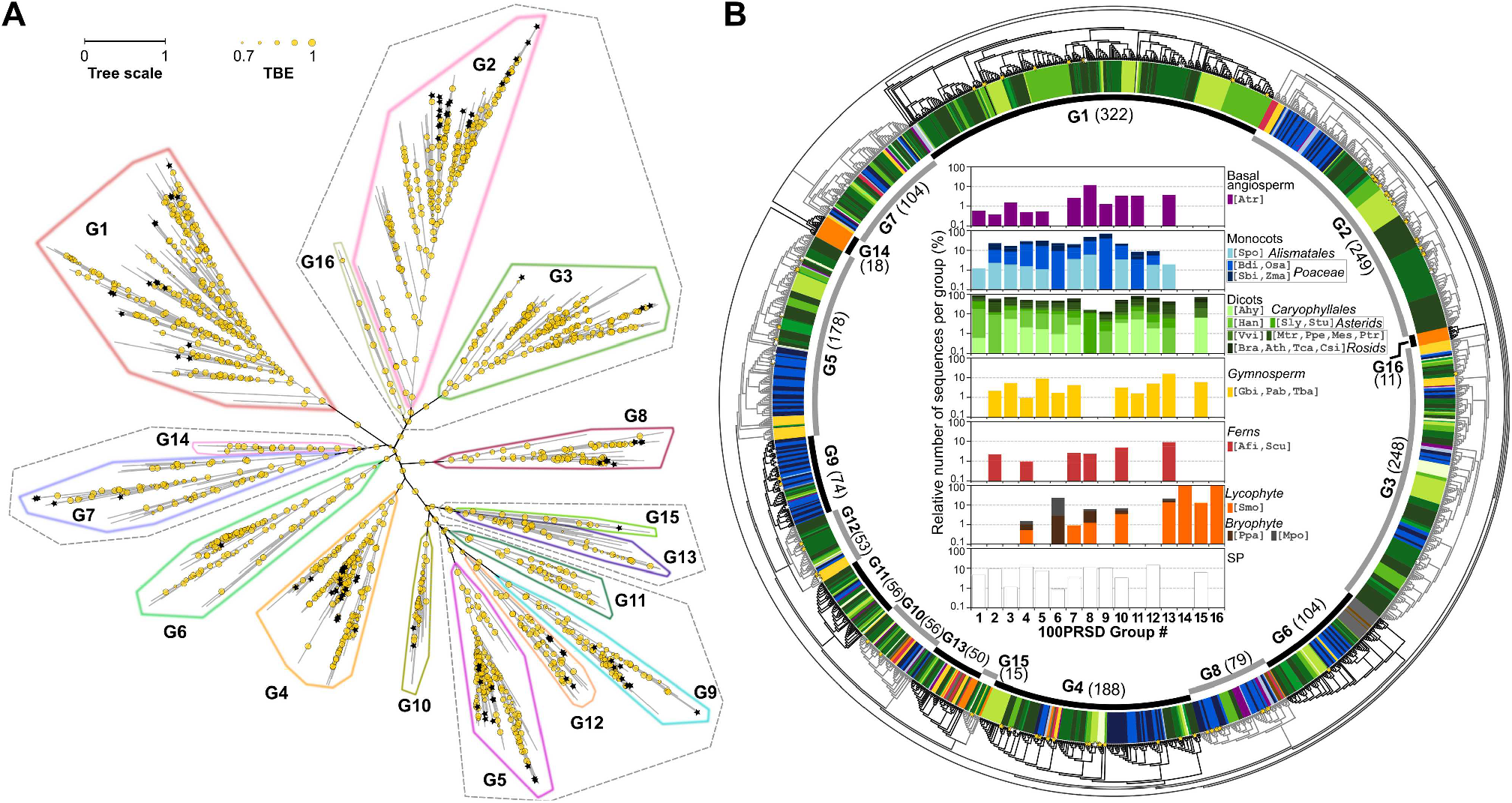
Maximum Likelihood *Phylogeny and HMMERCTTER clustering of land plant BAHD homologues*. (A) Radial phylogram representation of the phylogenetic relationships. The tree scale bar indicates 1 amino acid substitution per site. The branch statistical support, represented by TBE values obtained for 1000 bootstraps, is shown by circles (branches with TBE value > 0.7 are shown, the diameter of the circle denotes TBE value as shown by its scale). The sixteen identified high similarity clusters are contoured and labeled. Stars denote SP entries. (B) Circular inverted dendrogram with emphasis on taxonomic distribution, stars denoting SP entries. Outer strip: color representation of taxonomic description for sequences on each branch (colors as denoted in the center legend). Inner strip: each high similarity cluster is labeled with cluster id, and total number of sequences in round brackets. Center: relative number of sequences per cluster (log scale) belonging to each taxonomic group and species (See Figure 1 for species three letter code) or the set of SP entries.

In order to ensure a complete training dataset, we classify the reminder of sequences (n=120) that were not recovered after they were removed by our specific scrutiny with Seqrutinator (except partial sequences). This results in 44 classified sequences, yielding a final dataset with 1805 sequences, including two bryophytes genes from *P. patens* and *M. polymorpha* (PpaHCT and MpoHCT, respectively) with hydroxycinnamoyl transferase (HCT) activity and complementing capacity of *A. thaliana hct* mutants [22]. Upon analysis of these sequences, corresponding indeed to PpaHCT and MpoHCT, we confirm they were removed by our automatic scrutiny because they have a large insert (144 and 117 residues, respectively) not conserved in other BAHD sequences (Supplementary Figure S1A). This insert correspond to a highly-disordered, solvent-exposed loop in PpaHCT (Supplementary Figure S1B). Interestingly, removal of this insert in PpaHCT has only minor impact on activity and substrate selectivity [2]. As a result, G4 has now sequences from all species in the training dataset (with the exception of *G. biloba*). Another interesting update on taxonomic distribution following classification is the incorporation of one sequence from *S. moellendorfii* into G7.

#### Taxonomic distribution in the groups 100P&R-SD hierarchical clustering

The tree and its clustering are complex given the taxonomic distribution (**Figure 2B** and Supplementary Dataset S2). We first set out to describe the mayor evolutionary events taking into account that the evolution of the BAHD superfamily in land plants has been subject to birth and death evolution, which is particularly clear considering there is not a single cluster that has sequences from each species we investigated. If we consider that bryophytes and lycophytes represent ancient land plants taxa, half of the groups can be considered ancient and we envisage seven ancestral clades: G2+3+16 (n=508); G4 (n=188); G7+14 (n=122); G6 (n=104); G8 (n=79); G13+15 (n=65); and G10 (n=56) (**Figures 2A** and **2B**, Supplementary Dataset S2). Note also that the G1 (n=322) and G5+G9+G11+G12 (n=361) do not present sequences from ancestral taxa (extended also in this case to ferns). The G2+3+16 clade has two major clades with considerable taxonomic contribution, although lycophyte sequences are contributed by a cluster with eleven sequences from *S. moellendorfii* (corresponding to G16), which would represent ancestral sequences for clade G2+3+16 and invoke its origin in the lycophytes. G13 can be considered ancient, having sequences from lycophytes and bryophytes (as well as most other species, except for the Poacea), representing an example of early gene death. The G15 is small with its members scattered with one or two members in several but not all eudicots as well as *S. moellendorfii* and gymnosperm *P. abies*. This subfamily likely resulted from a duplication of a G13 sequence in a lycophyte ancestor but does not represent a core function. The G2+G3 has its origin in G14, which has 18 sequences in *S. moellendorfii* only. G2 likely does represent a core function given that each fern species on have at least one homologue.

G6, G8 and G10 all evolved from an ancestral Bryophyta homologue and all show clear events of gene death. G6 lacks a lycophyte and ferns homologues, and shows two homologues in *P. abies* but not in another gymnosperm. It also lacks homologues in *A. trichopoda* and *S. polyrhiza*. G8 lacks homologues in gymnosperms and certain other species, albeit that there is no pattern to suggest a single gene death event. G10 has homologues in all species except gymnosperms *T. baccata* and *G. biloba*. G4 also seems to have its origin in the lycophytes and only lacks a homologue in gymnosperm *G. biloba*. Groups 3, 5, 11 and 12 are more recent and correspond to spermatophytes albeit that G12 lacks an *A. trichopoda* sequence. Groups 9 and G1 are the most recently evolved groups only since they only have angiosperm sequences.

Interestingly, G1 is by far the largest group which is due to recent duplications in a number of eudicots, in particular *H. annuum* with a total of 57 G1 homologues. Indeed, only six sequences in G1 are not from eudicots (four sequences from *S. polyrhiza* and two from *A. trichopoda*). G9 has relatively many homologues from Poacea. Other species with relatively many homologues in a single group are G6 with 19 sequences in *M. polymorpha* and G14 and G16 that have 18 and 11 sequences, respectively, in *S. moellendorfii*. This high number of homologues in these lower plants are in contrast with the BAHD homologue distribution demonstrated by most higher plants.

Next, we checked the groups for the presence of SwissProt entries, since these are instrumental for the functional annotation of the various homologues. I we consider G14 and G16 as part of G2+G3 and G7 respectively, and that G13 and G15 also form a superclade, the only clade without any SP entry is G11. G12, G4, G8 and G9 appear as the best studied clades having 15.1, 10.4, 11.4 and 10.8% of their sequences as part of SP. The functional annotations of the SP entries from a single group are typically diverse. Also, given the large number of homologues per species of a number of groups, this indicates the identified clusters do not correspond to a single function but rather show further subfunctionalization.

### Characterization of BAHD homologues in groups 100P&R-SD indicates partial functional diversification

We used the clustered SwissProt entries to retrieve functional information of the homologues in the 100P&R-SD groups. This strategy can allow for a transitive functional annotation of the clusters. Out of the 113 SwissProt entries in the dataset (Supplementary Dataset S3A), we identified 85 of them with biochemical and/or physiological data in the literature (Supplementary Dataset S3 B). There is an extraordinary diversity of acceptor chemicals (93 not CoA-activated compounds) allowed by BAHD enzymes, including a variety of alcohols (both aliphatic and aromatic), diamines, glycosylated flavonoids and anthocyanidins, alkaloids, terpenoids and ciclohexanes (Supplementary Dataset S3 C). In order to find relationships among these compounds, we used 3D multidimensional scaling based on similarity indexes (such as Tanimoto index), and K-means clustering (**Fig 3** and Supplementary File S1).

**Figure 3:**
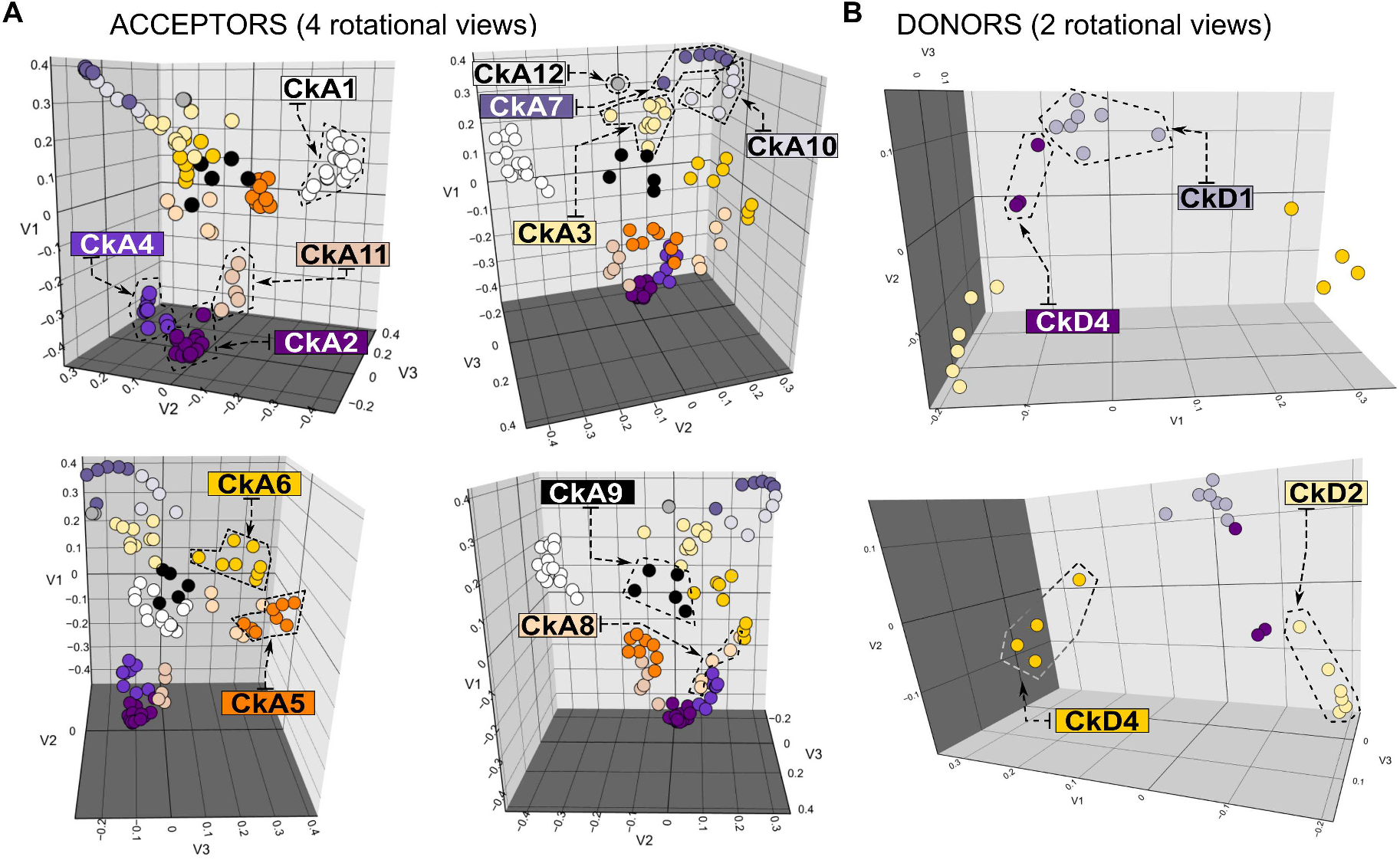
Characterized BAHD acceptors and donors clusterized by chemical features. 3D multidimensional scaling based on chemical properties resulted in a 3D plot, which was then clustered by K-means in 12 CkA clusters for acceptors (A) and 4 CkD clusters for donors (B). Clusters indicated by different colors. Four rotational views of the acceptor 3D plot and two views of the donor 3D plot are shown). For additional information see Supplementary Table S3.

For acceptor metabolites, acceptors were separated in 12 clusters (termed CkA1 to 12). CkA1 is an isolated cluster containing mainly aromatic alcohols (e.g., hydroxycinammoyl alcohols), plus a number of diverse aromatic primary amino compounds and hydroxyphenyl lactates (**Fig 3** and Supplementary File S1). Several neighbor clusters include different glycosylated compounds: in CkA11 we found smaller compounds, CkA2 contain flavonoids, isoflavonoids and anthocyanins, whereas CkA4 includes highly decorated anthocyanins. Another set of closely related clusters contain aliphatic compounds: CkA3 group a number of short chain alcohols, glycerol and diamines, CkA7 includes alcohols from four to ten carbons, CkA10 contain mostly longer chain alcohols (12-16 C), plus the polyamine spermidine and 15-hydroxypentadecanoate (a hydroxylated fatty acid), whereas CkA12 contain two isomers of a saturated 6 C alcohol. CkA5 includes a number of complex alkaloids (e.g., 17-O-deacetylvindoline). CkA6 clusters a number of *a priori* dissimilar compounds (brassinosteorids, anthocyanidins, 1,2-dyacylglycerol, diterpenoid and alkaloids. Another eclectic group is CkA8, including three diterpenoids (baccatins) and two differentially decorated acyl-sucroses. Finally, CkA9 includes quinate and shikimate, glucose, malate and methanol. Concerning donors, we have identified four CkD clusters grouping the 20 CoA-activated chemicals processed by the characterized BAHD enzymes. In CkD1, we found donors with aliphatic moieties (from two to six carbons, including acetyl-CoA) and dicarboxylates such as malate-CoA. Most closely related to this cluster is CkD4, which includes aromatic activated compounds such as benzoyl- and anthranoyl-CoA, plus tygloyl-CoA. CkD4 includes mostly (hydroxy)cinnamyl-activated compounds, such as coumaroyl- and caffeoyl-CoA. Finally, the cluster CkD3 includes CoA-activated long (C16-18) fatty acids and dodecanoyl-CoA.

We then cross referenced the instances in which acceptor and donor metabolites have been identified in biochemical and physiological studies for the BAHD enzymes annotated in the 100P&R-SD groups (Acceptors: **Fig 3A** and **Table 1**, Donors: **Fig 3B** and **Table 2**). In this sense, characterized BAHD enzymes in G1 and G2 can process acceptors from several distant CkAs, suggesting a plasticity towards chemically diverse metabolites (note, however, that acceptors in G2 correspond mainly to glycosilated metabolites). However, enzymes from G1 use donors from CkD1 (mainly acetyl CoA), and enzymes in G2 use donors from CkD2 as donors (with malonyl-CoA as preferred substrate) and from CkD2. In G4, there are 19 enzymes biochemically characterized. The acceptors in CkA9 are used mostly by these enzymes, particularly shikimate and quinate. However, there are used also chemicals from CkA1, CkA3 and CkA10. In terms of donors, those from CkD2 are vastly more used, particularly Coumaroyl-, Caffeoyl- and Feruloyl-CoA. These results suggest that enzymes in G4 is mainly associated to the HCT/HQT function, which typically involves Coumaroyl- or Caffeoyl-CoA to acceptors shikimate and quinate. Only one instance of Benzoyl-CoA (CkD4) is observed. Note that the enzyme using this donor processes anthranilate (CkA1) as acceptors (Supplementary Dataset S3 B). There is also an example of an enzyme using 5-hydroxyanthranilate, which may be involved in the biosynthesis of phytoalexins, and two enzymes in G4 that can process hydroxyphenyl lactates (also from CkA1) with Coumaroyl-CoA as donor, for the biosynthesis of rosmarinic acid characterized in Lamiaceae. However, rosmarinic acid (with a potential role in plant defense) has also been identified in land plants from a broad taxonomic range. Finally, we also identified an enzyme using spermidine (CkA10) as acceptor, shown as important in the synthesis of phenolamides (or hydroxycinnamoyl amides) in pollen of *A. thaliana*.

**Table 1:**
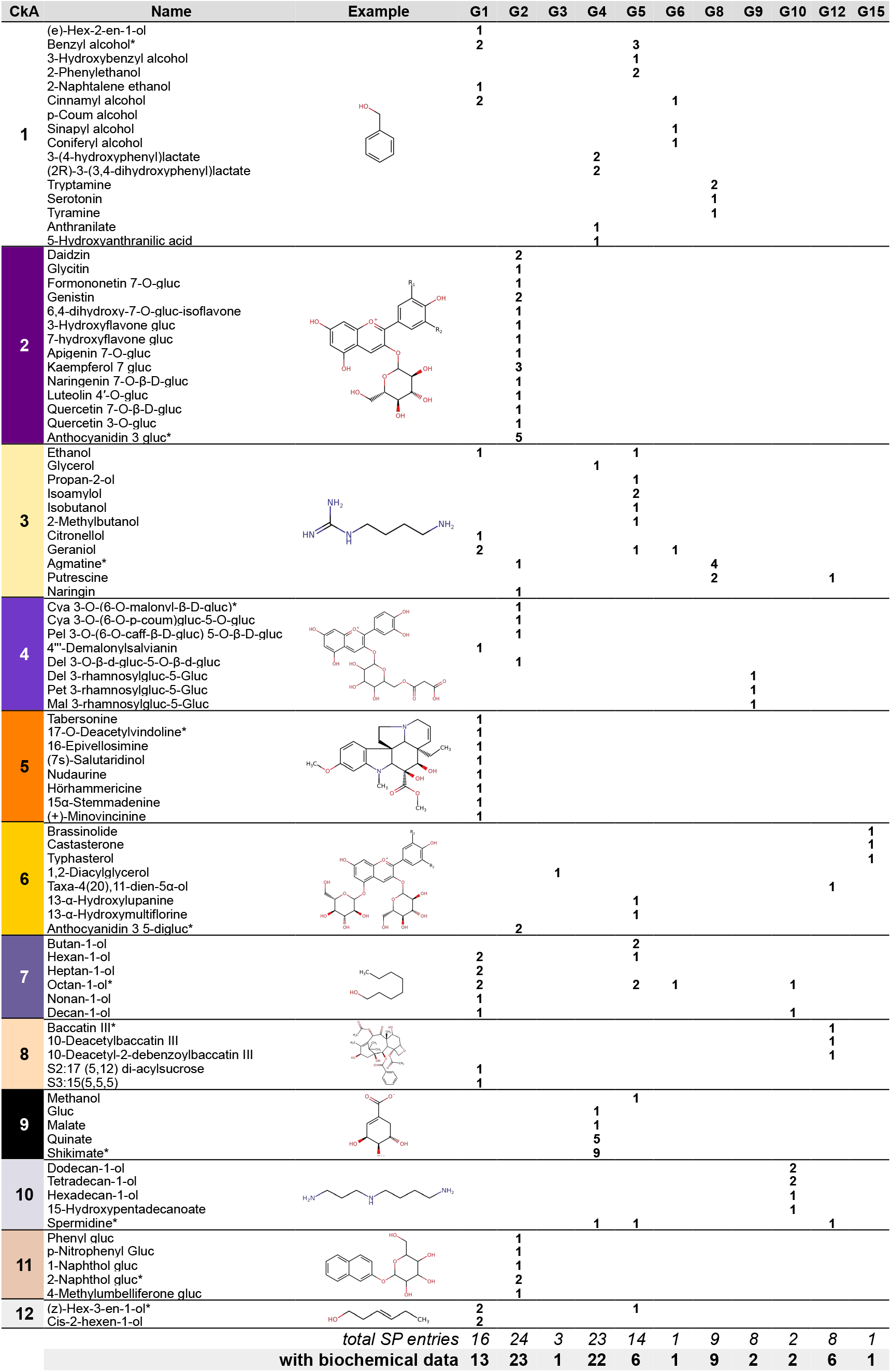
Cross data of clustered acceptors processed by BAHD enzymes in 100P&R-SD groups. Acceptors, shown in rows by clusters CkA1 to 12 (see also Figure 3). In the columns, count of instances in which a metabolite is processed by a characterized enzyme in each of the groups 100P&R-SD. Amount of SP entries per group are shown. Chemical structure of representative metabolites in CkA (indicated by *) shown. Abbreviations: Caff, Caffeoyl; Coum, Coumaroyl; Cya, Cyanidin; Gluc, Glucose; Mal, Malvidin; Pel, Pelargonidin; Pet, Petunidin.

**Table 2:**
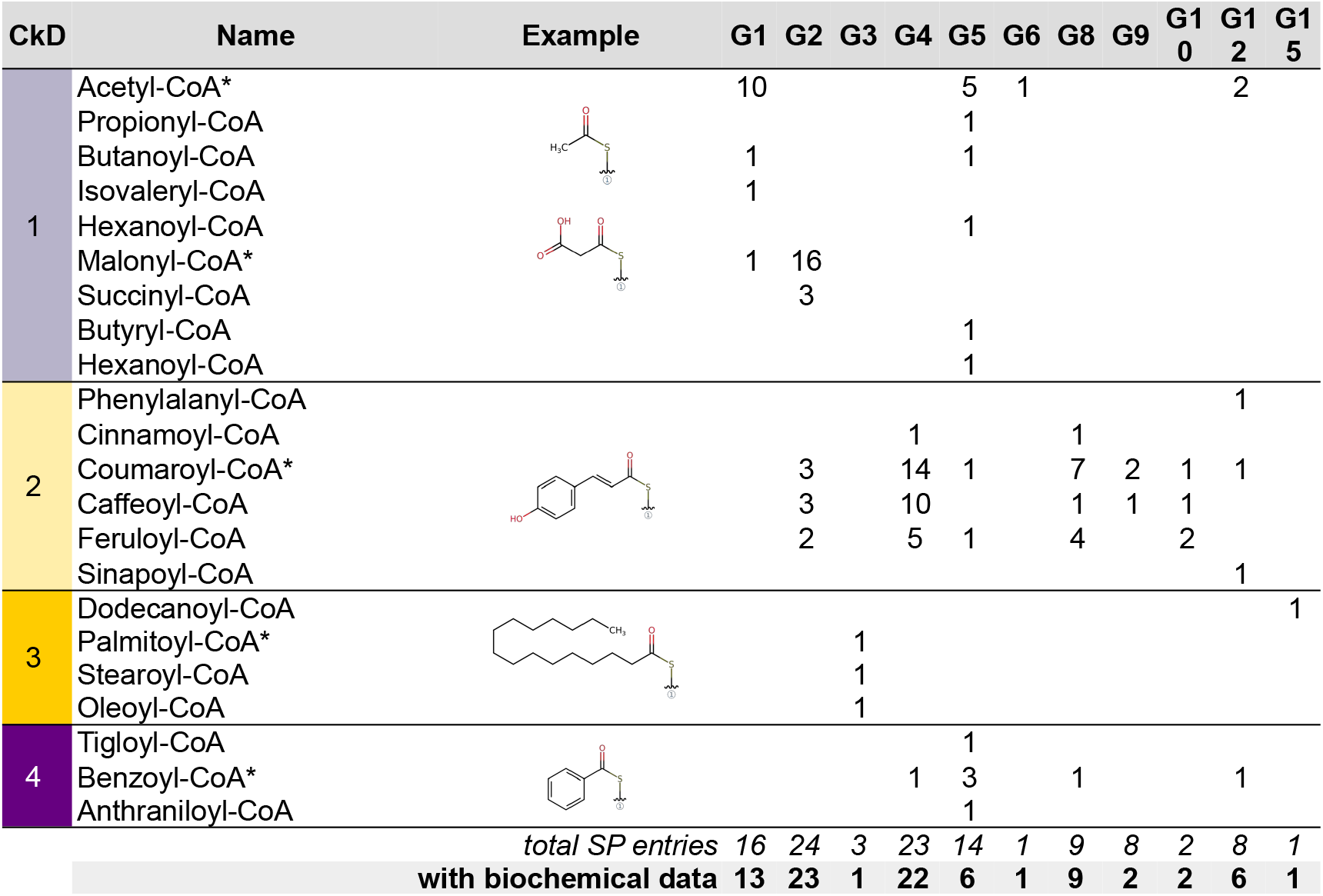
Cross data of clustered donors processed by BAHD enzymes in 100P&R-SD groups. Acceptors, shown in rows by clusters CkD1 to 4 (see also Figure 3). In the columns, count of instances in which a metabolite is processed by a characterized enzyme in each of the groups 100P&R-SD. Amount of SP entries per group are shown. Chemical structure of representative metabolites in CkD (indicated by *) shown. Note that Coenzyme A moiety is condensed. Abbreviations: CoA, Coenzyme A.

For G5 we observe a broad selectivity both towards acceptors (CkA1, 3, 6, 7, 9, 10 and 12) and donors (CkD1, 2 and 4). Note, however, that the majority of the acceptor compounds correspond to either aliphatic or aromatic primary alcohols, which are associated to the formation of volatile esters, typically by the transfer of CoA-activated aliphatic acids (mainly acetyl-CoA) or Benzoyl-CoA.

For G10, there are two SP entries biochemically characterized. Both enzymes, from *A. thaliana*, can process acceptors from closely related CkA7 and CkA10 groups, whereas they can process donors from CkD2. The role assigned to this activity is the synthesis of alkyl hydroxycinnamates, typically required for suberin formation. The fact that all taxonomic groups are represented in G10 with few copies per proteome suggest that the activity mediated by members of this subfamily correspond to a very conserved function across land plants.

For G12 (6 characterized enzymes, the acceptors processed belong to quite different chemical groups (CkA3, 6, 8 and 10), as well as the donors (CkD1, 2 and 4). There is, however, a potential link between taxonomic and functional signals that can shed light into this. Four out of the six characterized enzymes are from gymnosperms (specifically from *Taxus* genus), and they are all involved in different steps of taxol biosynthesis (Supplementary Dataset S3). The remaining characterized enzymes (O80467 and Q8GYW8 from *A. thaliana*) can be involved in the phenolamides synthesis based on the substrates they can use [23].

Finally, G15 has one characterized entry from SP. The enzyme can process brassinosteriods as acceptors (CkA6), and as donor it can utilize dodecanoyl-CoA (CkD3). The biological role associated to this enzyme is the modulation by inactivation of brassinosteroids in *A. thaliana*. Brassinosteroids are conserved across land plants [23]. However, the scattered taxonomic distribution in this group suggest a role conserved in some taxa, such as lycophytes, gymnosperms and some eudicots (typically Malvids, except from fabids *M. truncatula* and *P. trichocarpa*).

Taken together, these results show that the functional diversification of land plants, in terms of the chemical properties of the acceptors and donors they can process, is somehow driven by their phylogenetic relationships. However, this is not always sufficient to explain the functional diversification of the different groups, as other evolutionary processes may be involved in the diversification for this complex gene family.

### Intra-family sequence identity varies greatly BAHD families

In this sense, we analyzed percentage of identity (ID%) between all sequences in the dataset (**Fig 4**), including internal comparison (sequences of a group versus all sequences in the same group) and external comparisons (all sequences of a group versus all sequences another group). As general observations for internal comparisons, we compute ID% averages of ∼35% for G1, G2 and G7, slightly higher for G4 and G9 (∼45%), and highest for G10 with 60.5%. All internal distributions exhibit right-skewness at various degrees, ranging from a value of 25.5 in G1 to 0.4 in G10. In addition, Kurtosis scores suggest that all distributions have a long tail, except for G10 ID% distribution (which is in accordance also with the upper bound values, ranging from 45.3 in G1 to 90.7 in G10). As expected, there is a trend towards higher ID% in the internal analysis when contrasted to the external comparisons, although in general the highest density of points and the median in the internal comparison are still largely overlapping with the external comparisons plots for most groups (except G9 and G10).

**Figure 4:**
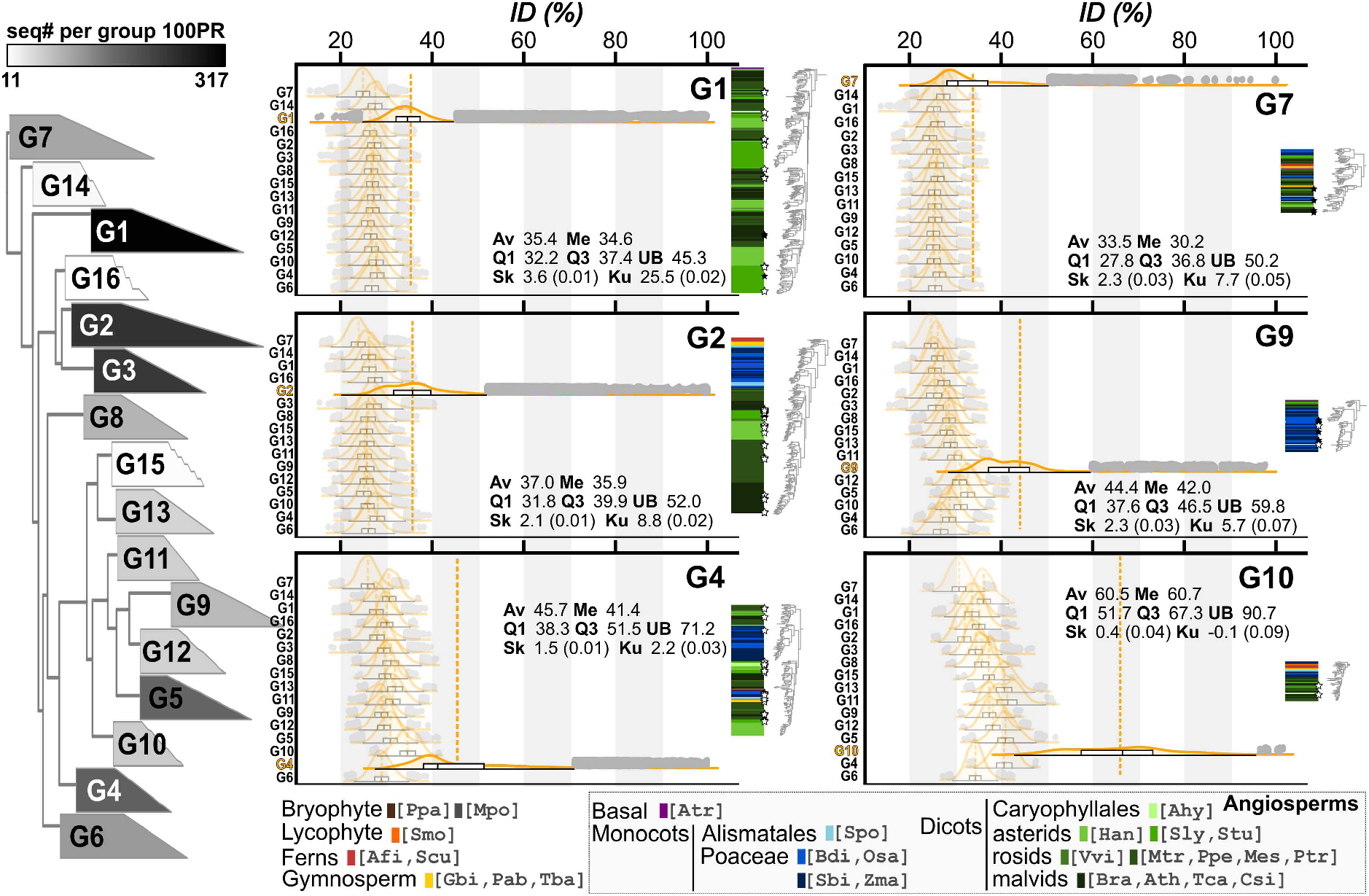
All-to-all sequence comparison of identity percentage. Distribution of percentage identity among all sequences in one group 100P&R-SD (particularly G1, G2, G4, G7, G9 and G10) compared to all sequences in each group. A schematic BAHD tree is shown to the left. Groups 100P&R-SD are collapsed, and group size is indicated by the color according to the gray scale shown above. Distributions plots are shown as split violins with its corresponding boxplot. Comparisons of sequences in one group with themselves (internal comparison) are shown with maximum opacity. Dashed lines, distribution average (extended line is average of the internal comparison). Outliers points are also shown. The order of the distribution plots correspond to the order of the groups on the schematic tree. To the right, a summary of the internal topology of each group, with the corresponding taxonomic representation of each leaf. Statistical values for internal comparisons are shown: Av, Average; Q1, first quartile; Me, Median; Q3, third quartile; UB, upper bound; Sk, Skewness (Standard Error); Ku, Kurtosis (Standard Error). SP entries are shown with stars (empty stars, characterized enzymes).

In the large groups G1 and G2, as in G7, the average ID% of the in internal comparisons partially overlaps with the tails of distribution plots of external comparisons. The trend is similar for these groups, even though G1 is taxonomically dedicated (almost all eudicot species) and present a broad range of biochemical activities, whereas G2 and G7 present most taxonomic groups and characterized activities are more conserved. One shared feature of G1 and G2 is the high number of copies per proteome typically observed (Supplementary Dataset S2).

As we already mentioned, there is a variable number of sequences per proteome in G4 (Supplementary Dataset S2) and a broad taxonomic representation. Note, however, that the internal topology of G4 suggests that there are subclades with differential taxonomic representation (**Fig 2B, Fig 4** and Supplementary Fig S3). One of the subclades presents all taxa represented (with low copy number per taxon), and it includes most of the characterized SP entries. The other two sublcades are taxonomically dedicated (one is all eudicots, and the other all monocots), with higher copy number per proteome. The former presents the highest internal ID% (Supplementary Fig S3). Therefore, the group shows mixed functional and taxonomical signals which may impact on the distribution of ID% towards lower values.

Finally, G9 and G10 are both medium/small size. In both cases, characterized activities in these groups correspond to conserved functions (phenolamides and suberin biosynthesis, respectively). However, for G9 copy number per proteome is high, particularly in monocots species, whereas that G10 presents a broad taxon representation an low copy number per proteome. Remarkably, the ID% trends towards higher values in G10, whereas G9 shows distribution towards lower ID% values.

ID% distributions in these scenarios suggest that multiple copies por genome are presumably subjected to different evolutionary rates, and contribute differently to functional signals, although.

### The expanded land plants BAHDome from additional 191 complete proteomes allows the identification of potential functional families in Eudicots and Monocots

In order to increase the amount of information, we created an expanded land plant BAHD dataset. We first performed a data mining in additional 191 land plant proteomes, resulting in a target dataset with 14267 sequences (Supplementary Dataset S4). Of these, 13799 sequences (96.7%) are classified into the training set, resulting in a BAHDome with 15607 sequences. With the exception of G14 and G16, all groups classified new BAHD homologues. The groups show a growth of 11.5 times (G11) to 6.1 times (G13), representing the ratio of sequences after and before classification. Since the target is 7.9 times larger than the training, this suggests that both training and target datasets are taxonomically balanced at some extent. In average, 96.8% sequences per species are classified (s.d. 8.3%). With the exception of *Fragaria* x *ananassa* dataset (for which only 6.4% of the sequences were classified), more than 62.3% of the BAHD sequences per species are classified (for 102 species, this number is 100%).

In terms of taxonomical distribution of sequences classified in each group, results agree with the trends observed in the training dataset. For example, of the 3004 sequences classified in G1, 2973 are from eudicots, being all orders represented (except from Proteales). The rest of the sequences belong to species from: i) orders Laurales and Magnoliales, closely related to eudicots (7 and 11 sequences respectively), ii) order Amborellales (2 sequences, from training), iii) more strikingly, 6 sequences from *Ananas comosus*, belonging to Poales order and (together with the four sequences from *S. polyrhiza* from the training), the only monocots sequences found in G1. Other groups with strong biased taxonomic representation are G8 (absent in gymnosperms and poorly represented or absent in eudicots) and G15 (scattered in some Angiosperms, most rosids, monocots, some Gymnosperms and Selaginellales, typically with one or two copies per species). In contrast, only 9 species do not classify sequences into G4 (different species scattered across different taxonomic groups) and into G3 (inferior plants not represented: ferns, Lycophytes, Marchantiophyta and Bryophyta). Other examples of groups with broad taxonomic representation are G10 (199 species from all groups); G2 (195 from all groups, including ferns but not Lycophytes, Marchantiophyta and Bryophyta); and G11, G5 and G6 with 195, 194 and 191 species represented (respectively), typically with low to no sequences from lower plants (with the exception of G6 with sequences from Marchantiophyta and Bryophyta). Note that in many cases, the BAHDome of species without sequences in a group with broad taxonomical distribution are often small compared to closely related species, suggesting low quality or incomplete proteomes. For example, *Ipomea trifida* lacks of homologues in G4. However, this species has a total of 51 BAHD homologues, which is low when compared to *I. nil* and *I. triloba*, with 107 and 82 homologues respectively. Similar observations are made for species lacking homologues in G4: *Corchorus capsularis* in Malvaceae, *Raphanus raphanistrum* vs *R. sativus, Corylus avellana* in Betulaceae, *Fragaria* x *ananassa* and *F. orientalis* vs other *Fragaria, Eragrostis tef* vs Poaceae and *Ginkgo biloba* vs other Gymnosperms.

The reconstruction of the BAHDome phylogeny with training plus classified target sequences show a topology almost identical to the topology observed for the training BAHDome (**Figure 5A**). The main observed difference is that sequences in G12 are distributed among two closely related but separated subclades, where the basal subclade contains all the sequences from Gymnosperms (including sequences from SP.

**Figure 5:**
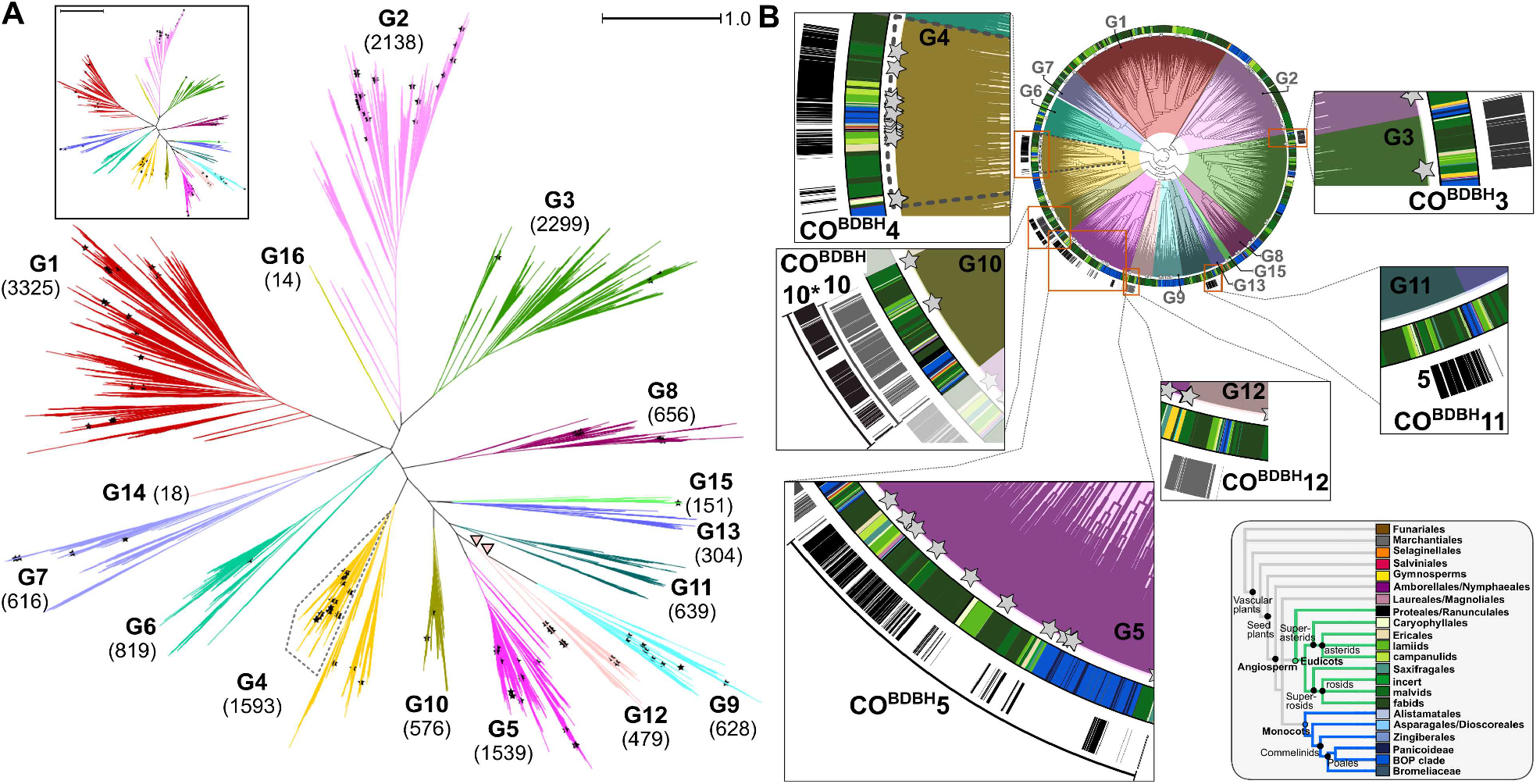
Phylogenetic reconstruction of BAHDome after target classification and identification of potential functional families. (A) BAHD sequences classified from target plus sequences from training were used to reconstruct a maximum likelihood phylogeny. Sequences are colored according their group identity after classification (in brackets, total number of sequences after classification). For comparison, the training tree with the corresponding groups is shown (inset). Pink triangles denote the origin of two branches defining G12. Scale line in both trees correspond to a distance = 1. (B) The same tree shown indicating sequences taxonomy, colored in the first inner ring according to the central panel. In addition, we show sequences detected as potential clusters of orthologues by bidirectional best BLAST hit (CO^BDBH^) using Eudicots and Monocots sets organized at order level. For both A and B, dotted line in G4 indicates sequences corresponding to G4’s sequences with wide taxonomic representation; stars are used to show SP entries.

In addition, we aimed to identify potential clusters of orthologues in the complete BAHDome using the bidirectional best BLAST hit strategy, which can be associated to functional families (see Methods for details). Because of the great diversity of sequences, both taxonomically and functionally, we limited the dataset to the Eudicots and Monocots sequences, organized at the hierarchic Order level. Hence, the dataset used contains 15411 sequences in 23 orders (we excluded Asparagales, Oxalidales and Zingiberales because of their low homologue number comparing to related orders). Another limitation we included was a threshold of 50% ID among hits, considering the typical large variation of BAHD sequences previously observed. As result, we have identified six potential cluster of orthologues by BDBH (CO^BDBH^, **Figure 5B** and Supplementary Dataset S5), including a total of 2016 sequences (13.08% of the sequences in the dataset). The largest is CO^BDBH^5, with 549 sequences, all identified to G5. Two characterized SP entries match to this CO^BDBH^, Q5I2Q5 and Q8GT20, both using aromatic alcohols as acceptors and at least Acetyl-CoA as donor (see also Supplementary Dataset S2 B). Note that typically most species are represented among orders, with the noteworthy exception of Brassicales, for which only 2 of 26 species are represented (Supplementary Dataset S5). The second largest cluster, CO^BDBH^4, contains 477 sequences, matching to G4 (and more specifically, the subclade in G4 corresponding to the complete taxonomical distribution (Supplementary Fig S3). A total of 10 SP entries were identified in CO^BDBH^4, 8 of them characterized biochemically (7 of them using shikimate/quinate as acceptor and all using Coumaroyl-, Caffeoyl- or Caffeoyl-CoA). In most orders, a high percentage of species are represented in CO^BDBH^4, except for Caryophyllales (20%) and Lamiales, Sapindales and Rosales (50, 55.6 and 56.5% respectively). Next, CO^BDBH^10 includes 401 sequences corresponding to G10, including one characterized SP entry (Q94CD1). A few interesting observations are worth to mention. On one hand, we also run the detection of CO^BDBH^ using a wider taxonomical set including not only Eudicots and Monocots, but also orders from basal Angiosperms (Amborellales, Nymphaeales, Laurales and Magnoliales), Gymnosperms (Pinales), ferns (Salviniales) and Lycophytes (Sellaginellales). It results in the identification of a sole CO^BDBH^ with 429 sequences, which we termed as CO^BDBH^10* because all sequences are also from G10. On the other hand, typically at least 77% of the species per order are represented in both CO^BDBH^10 and CO^BDBH^10*. Finally, CO^BDBH^10 and CO^BDBH^10* share 364 (78.8%) of the sequences. The fourth CO^BDBH^3 we have identified in the Eudicots-Monocots set is smaller, with 240 sequences belonging to G3. Note that because of the total size of G3 after classification (2299 sequences), CO^BDBH^3 corresponds to a minor subclade in this group (**Fig 5B**). Interestingly, CO^BDBH^3 holds one of the three SP entries described in this group. This entry, Q9FF86, has been described as important in cuticle formation, using 1,2-diacylglycerol as acceptor and activated fatty acids (such as Palmitoyl-CoA) as donors. We also observe a rather high representation of species per order (at least 66.7% of the species, except by Gentianales and Myrtales with 50%). Finally, we identify additional CO^BDBH^11 and 12 (with 185 and 164 sequences respectively). In the different orders we observe a high species representation, except by Poales in CO^BDBH^11 (3.7% of the species) and Caryophyllales in CO^BDBH^12 (40%). Note that none of the 6 SP entries in G12 that are characterized belong to CO^BDBH^12 (explained because five of them belong to Cupressales order, which is not in the analyzed set because belongs to Gymnosperms).

These results demonstrate the complexity of the land plant BAHDome, and that this complexity introduces a diversity that reflects in only 13.08% of the sequences in Eudicots likely corresponding to functional families.

### Specificity determining positions in BAHD families are typically not associated to the binding pocket

Next, we used the training dataset clusterized in 16 groups in order to identify specificity determining positions, SDPs. An SDP network (SN) with 33 SDPs was obtained (**Fig 5**) by combining data from estimated clustering determining positions, CDPs, and the mutual information (MI) between them. Note that SDPs are labeled according to the position in a reference protein, CcHCT (*C. canephora* HCT), which belongs to G4. It is important to indicate that we perform the identification of the SN using the training dataset, since a high quality MSA is instrumental to the identification of SDPs. Note, however, that we will analyze the conservation of the residues in the expanded BAHDome, as we describe next.

The resulting SN is depicted in **Fig 6A**. The score of the SN is significantly higher to the score of 1000 networks randomly generated with 38 nodes from the same dataset (z-score=70.05). The largest subnetwork identified with k-clique analysis has 11 nodes (subSN1), followed by a 10 nodes subnetwork (subSN2). Despite that several node pairs with high MI value belong to subSN1, note that the MICS node scores in this subnetwork are typically moderate to low values. The SDPs in subSN2 are connected by high, albeit slightly lower, MI values, but typically presenting higher MICS scores. Bear in mind, however, that for simplicity we have chosen to show only the highest quartile MI data; consequently the edges shown indicate already high MI z-scores.

**Figure 6:**
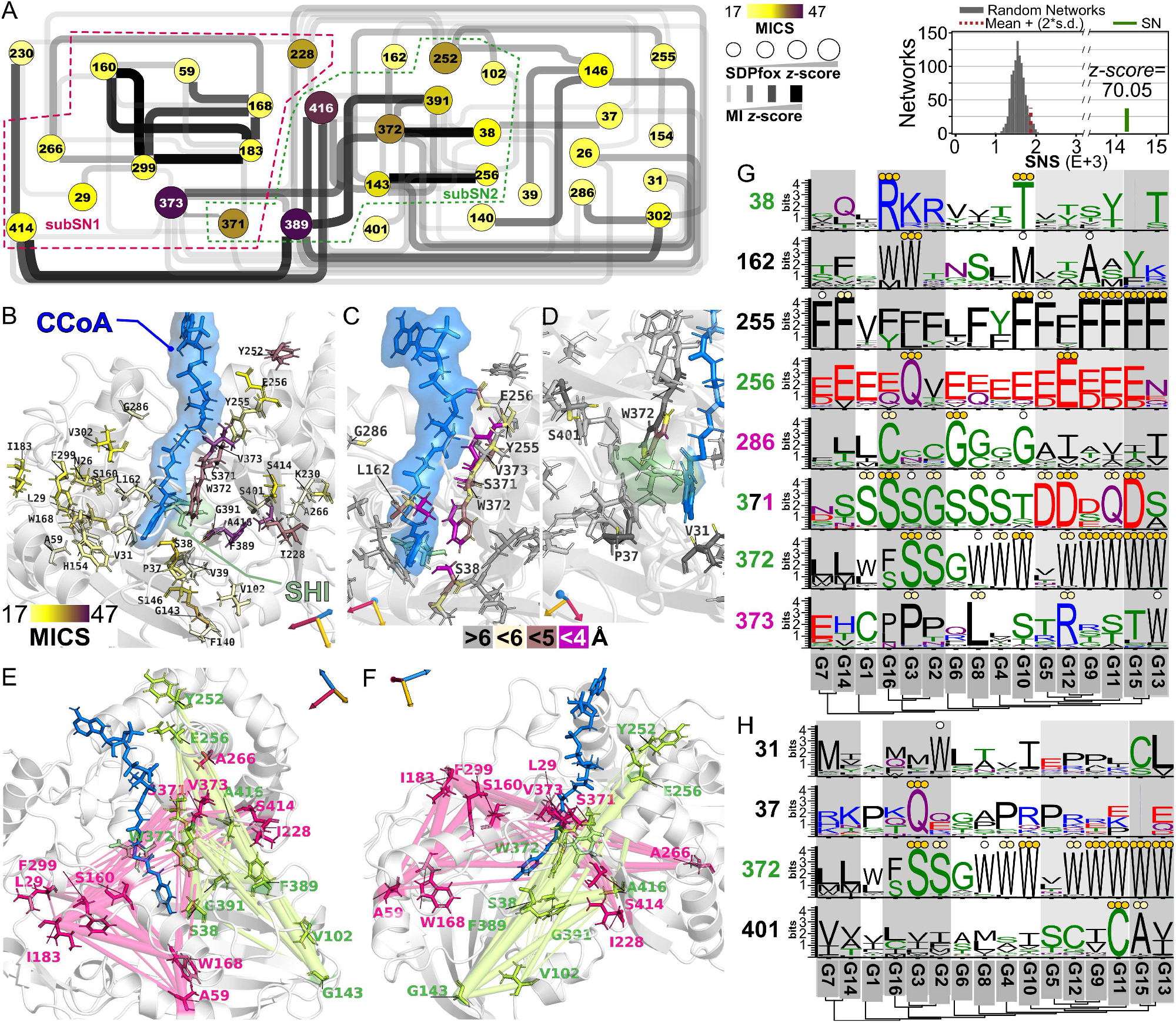
Identification of SDPs among BAHD subfamilies. Based on the clustered training set, an 33 SDPs network was identified. (A) SDP Network (SN). Nodes are SDP, colored according their MICS following the scale shown. Node diameter indicates value of SDPfox z-score (diameter increases for increasing z-score). Edges indicate the MI (color and width indicates MI value, as shown in the scale). For simplicity, only edges with MI > 10.7 (highest quartile) are shown. Two major subSNs are demarcated with colored dashed lines. Right: the SN score (SNS) is significantly higher when compared to 33-node randomly generated networks (n = 1000). (B) B to F: SDPs mapped in the structure of the reference CcHCT (Uniprot id A4KZE4) with donor coumaroyl-CoA (CCoA) and acceptor shikimate (SHI). Rotational coordinates indicated with 3D axes. In all cases, cartoon backbone is shown. (B) all SDPs are shown and colored according to their MICS. C and D: SDPs with atoms located at least at 6 Å from the donor (C) or acceptor (D) are labeled and colored according to the scale shown. Other SDPs are shown with gray sticks. (E,F) Two rotational views showing SDPs from subSN 1 (pink) and 2 (green). MI connections between residues are shown in transparent cylinders, where a higher thickness indicate a higher MI z-score. G and H: logos of SDPs located at <6 Å from donor (G) and acceptor (H) for each family. Families are organized following their phylogenetic relationships. Positions correspond to CcHCT (A4ZKE4), a reference protein from G4. Coloring in SDP indicate positions from the subnetworks indicated in A. Positions with residue conserved in ≥85%, ≥90% or ≥95% of the sequences of the group are indicated with •, •• or •••, respectively.

A final remarkable observation is that 14 SDPs are conserved in >85% of the sequences from G10 (8 of them conserved in > 95% of the sequences). This is consistent with the high conservation degree among sequences in G10 (**Fig 4**). In the same line, only 3 SDPs are conserved in G7, and 4 SDPs in G4, all groups with a large internal sequence variability. In the case of G4, only 6 SDPs with high conservation degree are found.

The SDPs were mapped in the reference structure CcHCT. First, all SDPs are shown and colored according to their MICS value (**Fig 6B**). In addition, because SDPs may be *a priori* linked directly to substrate specificity among members in different BAHD subfamilies, we mapped SDPs with atoms located closely (≤ 6 Å) to either the donor (**Fig 6C**) or acceptor (**Fig 6D**). It is interesting to mention that only a handful of SDPs are identified here: SDPs corresponding to CcHCT residues 38, 162, 255, 256, 286, 371 and 373 have atoms proximal to donor, residues 31, 37 and 401 with atoms proximal to acceptor and SDP 372 have atoms proximal to both substrates. We also explored the homologous SDPs in 3D structures of reference SwissProt entries from other groups (see Supplementary Fig S4). Because the SP entries from other groups can accept a number of different donors and acceptors, it could be expected that the binding pocket would be rearranged, resulting in SDPs that get closer to the substrates. However, in most cases, the same positions corresponding to the SDPs and proximal to substrates on CcHCT are indeed also proximal to the CcHCT’s substrates when other reference proteins are structurally aligned to CcHCT. However, other SDPs can be proximal to substrates in only some BAHD families, such as position 167 and 169 in G1’s and G7’s reference, respectively. Conversely, some SDPs are close to substrates in reference proteins from only some of the reference proteins (for example position 31 and 286 from CcHCT). Finally, homologous residues to SDP 31, 37 371, 372 and 401 can have atoms with proximity to either or both substrates in different references. Together, these results suggest that there may be a rearrangement of the catalytic pocket among references from different groups, although it does not impact on SDPs identified by our approach. Next, we mapped the SDPs from the identified subSNs (two rotational views in **Figs 6E-F**). From subSN1, only SDPs 371 and 373 have atoms proximal to donor. In subSN2, SDPs 38, 256 and 371 are located close to donor, whereas SDP 372 has atoms located in the vicinity of both substrates. Note that the residues in the subSN2 are spatially located close in CcHCT’s structure, which can be seen as a an interconnected cloud towards the right of the substrates in **Fig 3E**. Unlike this, the SDPs in subSN1 are located at larger distances, however connected with high MI values.

Finally, we obtained the corresponding logos for the 33 SDPs identified in the SN but using the datasets after target classification, in order to evaluate residue conservation in the complete BAHDome (Supplementary Fig S5). With the exception of SDPs 31, 230, 373, 389, 414 and 416, the SDPs show conservation in ≥95% of the sequences in at least one of the families. Of the remaining SDPs, only SDPs 29, 37, 286, 370, 373, 391 and 400 do not show conserved residues in ≥95% of the sequences in at least one group. A number of diverse scenarios are found considering conservation levels and physicochemical properties of the SDPs. For example, SDP 143 is largely conserved in most groups, typically presenting Gly, being the most remarkable exception G3, with Asp in >95% of the sequences, and G2, showing non conserved residues at this position. Similarly, SDP 168 is conserved in at least >90% of sequences from 9 families each, mostly with the bulky Trp residue at this position. SDP 140 is mostly represented by hydrophobic residues, with high conservation (>90% of sequences) of Phe in G1, G7, G9, G10, G12, G13 and G14, L in G3 and more variable in other groups. Another example is SDP 26, which shows highly conserved (>85% of the sequences) Asn in distant families (G14, G6, G10 and G13), and the acidic residue Asp only in G5, whereas a rather high variability degree is observed in the rest of the families. Considering that it is likely that specificity for donors and acceptors is mediated by those residues located on the vicinity of the substrates, we focused the analysis of the logos of these SDPs (**Figs 6G** and **6H** for residues nearby donor and acceptor, respectively). Regarding these SDPs, only three of them have residues conserved in >85% of the sequences from at least half of the families (SDP 255 in 8 families, SDPs 371 and 372 in 10 families). SDP 255 is represented by Phe in most families (including those without residues conserved in >85% of the sequences), or other hydrophobic residues in G1, G4 and G6. SDP 371 presents either polar Ser (G1, G3, G8 and G16), acidic Asp in G5, G12 and G15, basic Gln in G11 and small non polar Gly in G2. Finally, SDP 372 is conserved as Trp in G4, G8, G9, G10, G11, G12, G13 and G15, and as Ser in G2 and G3. Note that unlike SDPs 255 and 37 which have atoms located nearby the donor, SDP 372 has atoms located nearby both substrates. Other SDPs located nearby the donor have residues conserved in >85% of the sequences in only few families, such as SDPs 38, 162, 256, 286, 373 (**Fig 6G**). More strikingly, this is the case for all SDPs with atoms in the vicinity of the acceptor, with the exception of SDP 372 (which, as already mentioned, has also atoms near the donor, **Fig 6H**). Another interesting observation is that SDPs associated to the acceptor do not belong to the major subSNs (again with the exception of SDP 372 which belongs to subSN2). Conversely, concerning SDPs located in the proximity to the donor, only 162 and 255 do not belong to either subSN.

### The identification of SDPs in a functional family HCT/HQT (G4-1)

HCT/HQT enzymes (included in G4) are among the most characterized enzymes in the BAHD superfamily. These enzymes are of wide interest because of their association to the chlorogenic acid (caffeoyl-quinate) pathway, and biosynthesis of lignin precursors. Note however that several of G4’s enzymes can utilize other hydroxycinnamoyl-CoA as donors, and shikimate is also widely used as acceptor. The original G4 dataset was subjected to an inception analysis, which resulted in 3 subgroups 100P&R-SD (Supplementary Figure S6A), corresponding to our previous observations (Supplementary Figure S3). Using the clusterized G4 as training, we performed the classification of a target dataset (including sequences from the scrutiny not recovered plus the 191 BAHD sequences from additional 191 proteomes) resulting in 1411 sequences from the target classified into the three subgroups. As a result, G4-1, G4-2 and G4-3 result in 801, 328 and 466 sequences, respectively (Supplementary Figure S6 B). The resulting largest subgroup G4-1 contains all taxa, G4-2 contains only monocots sequences, whereas G4-3 includes only eudicots sequences (Supplementary Figure S6B). Note also that 20 of 23 SP entries belong to G4-1. Therefore, sequences in G4-1 match with sequences in CO^BDBH^2 (**Figure 5B**). The only SP entry in G4-2 corresponds to an HCT from rice (Q5SMM6), with additional capacity of glycerol utilization. In G4-3 we identified two SP entries (B6D7P2 and O64470), which can distinctively use malate or spermidine as acceptors, respectively (Supplementary Dataset S3). Thus, we considered the subgroup G4-1 as HCT/HQT subfamily.

Then we performed a dedicated analysis to identify specificity determining positions (SDPs) in G4-1 using the training dataset. Because SDP identification is a comparative analysis and it is largely dependent on the datasets used, we sistematically compared the G4-1 dataset to each of the remaining sets, except for groups G14, G15 and G16, due to their small sizes (18, 14 and 11 sequences, respectively). Again, we utilized the training datasets because it results in more compact MSAs, resulting in a more reliable SDP identification. As such, we performed a total of 13 comparisons, including a comparison with the remainder of the G4 (sequences from subgroups G4-2/G4-3 combined). All resulting SNs are summarized in Supplementary Figure S7. In total, 168 SDPs were identified (Supplementary Dataset S6). The number of SDPs detected in the comparisons ranged from 12 (vs. the remainder of G4, G4%) to 36 (vs. G11). Seventy four of the 170 SDPs were identified in only one comparison, 40 in two comparisons, 22 in three comparisons, and 32 SDPs were identified in four or more comparisons. Of the latter, the most remarkable are SDPs 67, 152, 261, 304, 334 and 391 detected in seven comparisons, SDPs 23, 266, 270 and 286 in six comparisons and SDPs 35, 146, 160, 183, 267, 299, 331, 371 and 389 in five comparisons. Interestingly, SDPs 261, 266, 267 and 270 are located in the helix α21 of the reference sequence CcHCT (**Figs 7A** and **7B**). In addition, SDPs 389 and 391 are located in sheet β31 (which also harbors SDP 390, identified in three comparisons).

**Figure 7:**
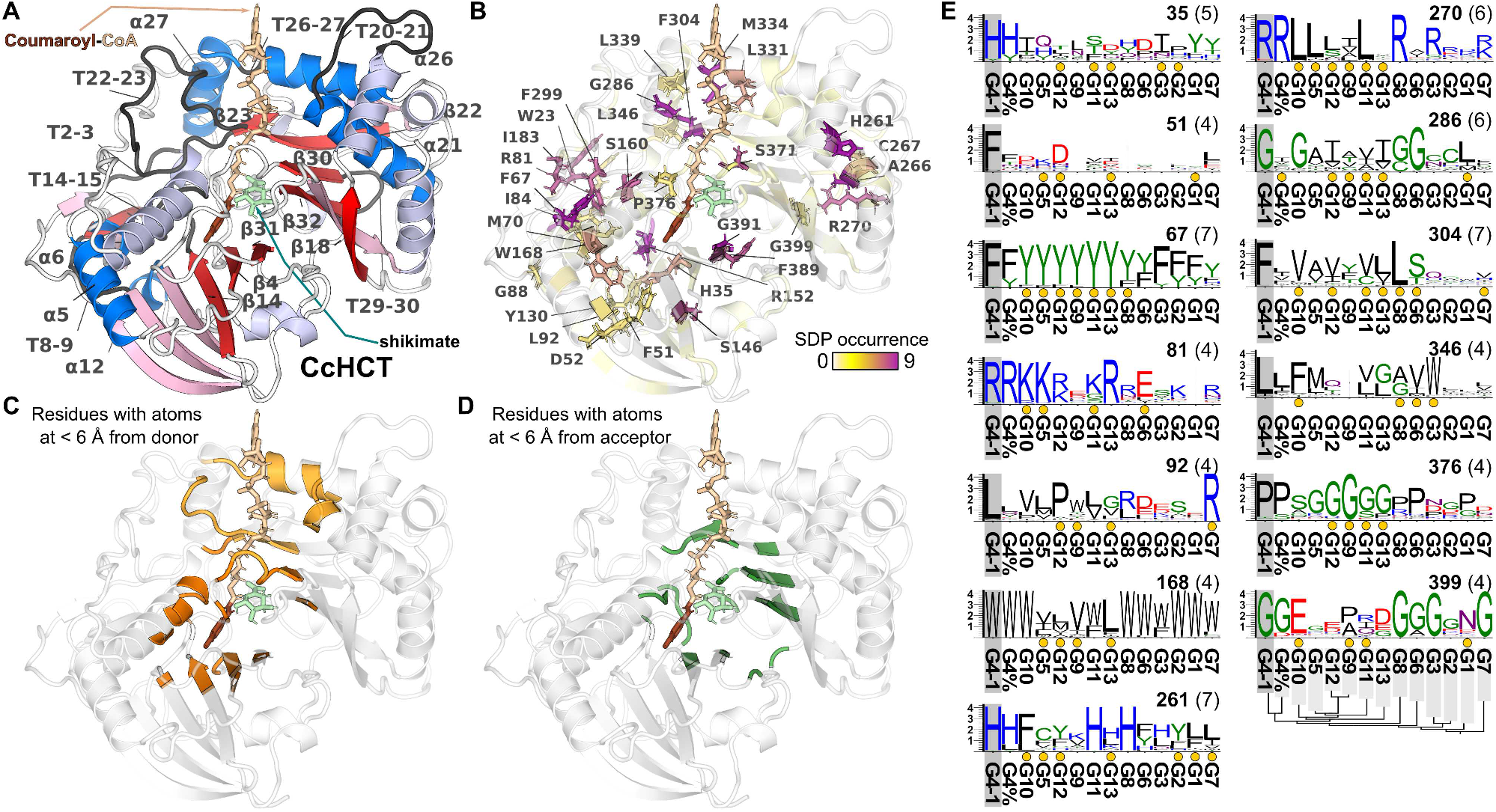
Identification and mapping of SDPs in HCT/HQT group. (A) Reference structure CcHCT with donor coumaroyl-CoA and acceptor shikimate. Secondary structures (alpha helixes, α and beta sheets, β) are colored in blue and red shades. Secondary structures with SDPs identified in at least four comparisons are shown in darker shade. Only these secondary structures are labeled, along with turns (T) fulfilling the same condition. Numbering correspond to their occurrence from the N-end. In donor, coumaroyl moiety is presented in a darker shade. (B) Identified SDPs are colored over the cartoon depiction using the scale shown. Sticks and labels are shown only for SDPs identified in at least 4 comparisons. Amino acid residue corresponding to the reference CcHCT is indicated in the SDP label. (C, D) CcHCT residues with atoms located at least at 6 Å from donor (C) and acceptor (D). Residues with proximity to the coumaroyl moiety in C are presented in a darker shade. Rotational axes for CcHCT coordinates shown in A. (E) Sequence logos of SDPs identified in at least four comparisons and conserved in ≥90% of G4-1’s sequences after target classification. SDPs are indicated in top right (in brackets the number of comparisons in which the SDP was found). Yellow dots indicate comparisons in which the position was actually identified as an SDP. The phylogenetic relationship among groups is shown. Logos x-axes values indicate bits. G4% indicates the remainder of G4, i.e. sequences in G4 not belonging to G4-1.

A total of 32 SDPs have atoms located near (≤ 6 Å) to the substrates (**Figs 7B, 7C** and **7D** and Supplementary Dataset S6). Regarding proximity to donor, SDPs 253, 255, 256, 282, 285, 286, 328, 331, 334, 371, 373 and 374 have atoms proximal to its CoA moiety, whereas SDPs 35, 38, 40, 150, 151, 162, 165, 375 and 390 are proximal to its Coumaroyl moiety. Conversely, SDPs 281, 362, 369 and 392 have atoms nearby the acceptor. Additionally, there are SDPs with proximity to both acceptor and either CoA (SDPs 283 and 304) or Coumaroyl (SDPs 36, 158, 370, 372 and 402) moieties of the donor. Note that only 6 SDPs with atoms proximal to substrates were found in ≥ 4 comparisons (SDPs 35, 286, 304, 331, 334 and 371).

Because the identification of SDPs depends on comparison of columns in different MSAs (i.e., for each comparison MSAs are built with the sequences from G4-1 and the sequences from the contrast group), SDPs can present different conservation degrees in the clusters compared. In order to analyze the complete BAHDome, the SDPs that we identified in the training dataset were used to investigate the conservation degree in the dataset after target classification, using comparative sequence logos (**Fig 7E** and Supplementary Figs S8 and S9). In this sense, several of the SDPs that were identified in more than comparisons are conserved in less than 90% of the G4-1 sequences (Supplementary Fig S8 and Supplementary Dataset S6). For example SDP 152, identified in 8 comparisons, is poorly conserved in G4-1 (Gln in 47% of the sequences, Supplementary Fig S9). Indeed, this position present highly conserved Asn in G3 (95%) and G10 (90%), and different degrees of conservation of Asn or Ser (ranging from 32% to 75%) in groups in which the position was identified as SDP (vs. G5, G9, G11, G1 and G7). However, it is noteworthy to mention that the reference sequence CcHCT has Arg in this position, an amino acid poorly represented in the rest of G4-1’s sequences. Along this line, other SDPs with residues poorly conserved in the sequences in G4-1, but highly conserved in other groups, can be mentioned. For example, SDPs 88 (with Asp conserved in G1 and G7), 266 (Arg in G5, G9 and G10 or Trp in G11) and 331 (Lys in G5, G9 and G12) are worth to mention (Supplementary Fig S7 and Supplementary Dataset S6). Other SDPs falling in this category are 52, 70, 183, 267, 299, 331 and 389.

Conversely, we can mention SDPs identified show a residue conserved in at least 90% of the sequences from G4-1’s, but not clearly conserved in any other group (**Fig 7E** and Supplementary Fig S9). In this sense, SDP 35 is mostly represented by His in G4-1 sequences, and mainly variable in the rest of the groups (except by the remainder of G4 in which His or Tyr can be found), whereas SDP 51 presents Phe in G4-1 but not conserved in any other group (the maximal is Asp in 67% of the sequences in G12). SDP 67 is largely conserved as Phe in G4-1, and consistently conserved as Tyr in the groups corresponding to comparisons that identify this SDP (**Fig 7E**). Conversely, SDP 168 is mainly conserved as Trp in most families, except on those in which this position is identified as an SDP. SDPs 81, 270, 376 and 399 show a conservation in most sequences of G4 (including G4-1 and also its remainder). In this sense, SDP 81 presents Arg in G4 and G13, Lys in G10 and G5 and Glu in G6. SDP 270 shows conservation of Arg in most sequences from G4, G8 and (at a lower extent), G3. However, it is replaced by aliphatic residues (mostly Leu) on several of the contrast families. Note that G4 presents the rigid Pro in G4, compared to predominance of the less restrictive Gly in several groups (as G12, G9, G11 and G13). SDP 304 shows high conservation of Phe in G4-1’s sequences, whereas mainly aliphatic residues, Ser or Cys is found in other groups. Finally, SDPs 261 and 286 are highly conserved in sequences from G4-1, less conserved in sequences from the remainder of G4, and residues with the same physico chemical properties conserved in sequences from some of the other groups.

A final noteworthy observation is that among the SDPs identified in at least four comparisons, four of them (SDP 129, 130, 389 and 390) do not present residues that are conserved in at least 85% of the sequences in any group (see Supplementary Figs S8 and S9).

Altogether, these results suggest that functional diversification of BAHD, taking HCT/HQT group as paradigm, depends on variability of residues that not necessarily are associated to the substrates binding pocket.

## DISCUSSION

In this work, we aimed to characterize the BAHD family in land plants, including information from a large number of complete proteomes from a broad taxonomical set. As such, employing a combination of sensitive (hmmsearch) and specific (Seqrutinator) datamining approach, we identify a total of 1805 sequences from 27 land plant proteomes and the characterized enzymes from SwissProt database (**Figures 1** and **2A**, and Supplementary Datasets S1 and S2). This training dataset was clustered into 16 groups, based upon the idea that the sequences on each monophyletic group can detect themselves with 100% precision and recall (100P&R-SD). In other words, the sequences on each group share more similarity among them to any other sequence from any other group. Furthermore, and following the same rule of 100P&R-SD, we classified a total of 13799 BAHD sequences from a target set from additional 191 complete proteomes, yielding a complete BAHDome with 15607 homologues (**Figure 5A**, Supplementary Datasets S4 and S7). Both training and complete BAHDome sets were used to understand different phylogenetic and functional traits of the BAHD acyltransferase superfamily. In general, the topology of the phylogenetic reconstruction generated (both with training and complete BAHDome sets) agree with others published elsewhere [21], [24]. These studies were dedicated to few species and, since they were mostly based on characterized BAHD enzymes, the complete BAHDome space was not extensively covered. In this sense, we detected a group G11 (including most Gymnosperm and Angiosperm species, Supplementary Dataset S4) that has no characterized SP entries and, to the best of our knowledge, no counterpart in other reconstructed BAHD phylogenies. As such, this G11 represents an unknown function BAHD family in superior land plants. In terms of homologue copy number, our results agree with the observation that during evolution of land plants, there was a consistent increase in the homologue copy number of genes in several protein families involved in specialized metabolism (including BAHD) [25]. This increment may be due to different processes of gene birth and death across different taxa, but also to massive genome duplication events. In this sense, we have identified 36 homologues in *Triticum urartu*, a diploid ancestor of the hexaploid modern wheat *T. aestivum* [26], of which we detected 461 BAHD homologues. Moreover, 205 and 277 homologues were found in tetraploid species *T. dicoccoides* and *T. turgidum* (Supplementary Dataset S4). A similar observation corresponds to the amount of homologues in the tetraploid *S. tuberosum* and its diploid wild relative, *S. commersonii* (97 and 54, respectively) [27].

The BAHD have an ancient origin in land plants, with functional homologues already described in bryophytes [22] and lycophytes [28]. Considering this, we identified a number of groups and superclades that can all be traced back to the families to these taxa (**Figure 2B** and Supplementary Dataset S2). As such, only G1 and major clade G5+G9+G11+G12 cannot be accounted as ancestral clades (lacking also ferns sequences). G1, which mostly includes multiple copies per species in Eudicots, represent a complex pattern of recent gene birth and duplication events. This could be related to the fact that characterized enzymes in G1 can process a range of diverse acceptors (**Figure 3, Table 1 and** Supplementary Dataset S3). Interestingly, the median value of global ID% among the sequences of G1 is 34.6%, with minimum value of 15% (**Figure 4**). As a counterpart, G10 is represented by all taxonomic groups, including all inferior land plants, with low copy number per species and the largest ID% average among the sequences in the group (**Figure 4 and** Supplementary Dataset S4). Besides, this is the only group that resulted in a group of orthologues when orders from all taxonomic groups were analyzed (**Figure 5B** and Supplementary Dataset S5). The characterized SP entries in G10 can conjugate CoA activated hydroxycinnamoyl acids to aliphatic alcohols [29], [30], which are key components of suberin. These acceptors belong to neighbor acceptor clusters (CkA7 and 10, **Fig 3** and Table 1). Since most likely suberin biosynthesis have evolved early before land colonization by plants [31], our findings suggest that BAHD homologues in G10 have played an essential role in the process and may represent the ancestral BAHD superfamily role. A similar observation can apply to the subfamily in G4-1, containing sequences with HCT/HQT activity, principally linked to transfer of hydroxycinnamoyl-moietes to either shikimate or quinate (CkA9 and CkD2, **Figure 3**, Table 1 and Supplementary Dataset S3). In this subgroup, there is also a high taxonomic representation and may represent an orthologues cluster (**Figure 5B**). However, compared to G10’s observation, average copy number per species is higher and intra-ID% is typically lower in G4 as whole (**Figure 4** and Supplementary Dataset S4), although the ID% in the subgroup 4-1 (which actually corresponds to a group of orthologues with HCT/HQT activity as shown in **Fig 5B**) is typically higher (Supplementary Figure S3). HCT/HQT activity is important for lignin formation, a key component of cell walls in vascular plants. However, even though mosses do not present lignin, it has been suggested that lignin-like precursors may occur in non-vascular land plants [32], [33]. Most of the characterized SP entries (19 of 21) in G4 belong to G4-1. Thus, whether G4-2 and G4-3 correspond to functional diversification specific to Monocots and Eudicots (respectively, see Supplementary Figs S3 and S6) is difficult to ascertain in lieu of the lack of biochemical data. In G4-2, we have identified a rice sequence (Q5SMM6) with a putative role in hydroxycinnamoyl-glyceride formation *in vitro* (although it can also form hydroxycinnamoyl-shikimate) [34]. Although hydroxycinnamoyl-glycerides may have a commercial importance as water soluble antioxidants, their physiological role is unknown. In G4-3, on the other hand, two SP entries are characterized: O64470 from *A. thaliana*, with a role in the synthesis of conjugates of hydroxycynnamoyl and spermidine in tapetum cells [34], and B6D7P2 from *Trifolium pratense*, involved in the biosynthesis of phaselic acid (using coumaroyl-CoA and malate as donor and acceptor, respectively) [35]. It has been recently suggested that the OH-acylation of aliphatic and aromatic alcohols may represent the ancestral set of acceptors for land plants BAHD family, as it most likely existed before land plants emergence [24]. Our results agree with this observation, and in addition offer the notion that CoA-activated hydroxycinnamic acids constitute the ancestral donors. Indeed, the subfamily of 4-coumarate lyase (4CL), responsible for the biosynthesis of hydroxycinnamic acid-CoA in an ATP dependent reaction [36], has been recently shown to have homologues in ancient land plants such as *P. patens* [37].

We also detected other important functions that can be associated to orthologues in Eudicots and monocots associated to groups G3, G5, G11 and G12 (**Figure 5B**). Although the sequences in CO^BDBH^3 correspond only to a subclade in the large G3, the characterized enzyme in this cluster of orthologues (*A. thaliana* Q9FF86) implicate a key role of this BAHD subfamily in cuticle formation [38], an adaptation that was instrumental to land colonization as it helps to avoid water loss, which biosynthetic genes have been identified in lycophytes [39]. Although this CO^BDBH^ was identified only for Eudicot-Monocot orders, note that (i) basal to the CO^BDBH^, a number of gymnosperm sequences are found and (ii), a Seleginella-exclusive group is found basal to G2+G3 (**Figures 2** and **5A**), implicating that indeed, the ancestral function for the G2+G3 may have been the cuticle formation with its origin in Lycophyte, although biochemical or physiological characterization will be required. Another interesting insight is that the reaction catalyzed by the enzyme in this CO^BDBH^ involves quite dissimilar acceptors and donors as the reactions in the other CO^BDBH^ mentioned so far (**Table 1**). As such, the donor is an activated fatty acid (CkD3) and the acceptor is diacylglycerol (CkA6, which in turn includes several dissimilar compounds). Regarding other CO^BDBH^s such as 5, 11 and 12, note that they are all associated to the major cluster G5+G9+G11+G12, which, as already mentioned, include exclusively Gymnosperm and Angiosperm species. Interestingly, characterized enzymes in CO^BDBH^5, Q5I2Q5 and Q8GT20, have been implicated in biosynthesis of volatile compounds [40], [41]. Note that this CO^BDBH^ covers G5 sequences in an scattered way. This suggest a multiplicity of duplication events in this group. Although volatile ester formation is not the only role for enzymes in G5, because roles of these compounds are diverse and related to several interactions (e.g., competition, mutualism or defense [42]) we can envisage an adaptative role of the generation of these compounds, which could explain the large amount of homologues per species in the group. Hence, the roles for G5 enzymes (and potentially the major cluster) can be associated to physiological and ecological functions acquired more recently by land plants. In this sense, reactions characterized in G12 are linked to the biosynthesis of the diterpene taxol in yew trees (genus *Taxus*), which acts in plant defense as an anti-fungal [43]. Surprinsingly, it is known that yew endophytic colonizing fungi species (such as *Paraconiothyrium*) can also produce taxol and work in a mutualistic manner with the tree’s cells to avoid growth of parasitic fungi, suggesting a long time co-evolutionary history of the taxol biosynthesis between yew tree and *Paraconiothyrium*, which have been suggested to act as actual immune cells in some gymnosperms lineages [43].

In the case of the large G1 and G2, the large number of sequences can be explain in association to the large space of compounds these enzymes can process. In G1, we concluded that these enzymes are largely associated to incorporation of acyl groups to a number of different metabolites, mostly in Eudicots (**Figures 2** and **3, Table 3** and Supplementary Dataset S3). Most alkaloids in the dataset are processed by enzymes in G1. Alkaloids come in an impressive variability in structure, size and functions [42]. Consequently, a multiplicity of BAHD isoforms may be required in order to process this large chemical space. Similar conclusions can be drawn from G2 enzymes, which characterized enzymes are largely involved in decoration of anthocyanidins or other glycosilated flavonoids (**Table 3** and Supplementary Dataset S3B), also associated to multiple biological roles and covering a large chemical space [42].

Finally, in G7 we have identified 4 SP entries (termed ECERIFERUM and ECERIFERUM-like genes), but they have not been biochemically characterized. Nonetheless, it is important to mention that these enzymes (3 from *A. thaliana* and one from maize) have been associated to cuticular waxes formation [11], [44]. The role of these enzymes has been associated to elongation of very long fatty acid chains. Although the exact mechanism of this process is yet to be elucidated, several models have been proposed [11]. In this study, we found that all the taxa are represented in G7 (indeed, 208 species are represented in the complete BAHDome, Supplementary Dataset S4), suggesting a very important function conserved across land plants. However, there are several important differences that support the notion of a potentially different mechanism involving BAHD enzymes in G7. First, the His residue in the HxxxD motif is less conserved as compared to other groups (Supplementary Fig S2), with the exception of G14. Another difference, already reported but confirmed in our dataset, is that the motif DFGWG is completely lost in G7 (Supplementary Fig S2). Finally, these enzymes may be associated to ER membrane [45], [46], unlike all the other soluble BAHD enzymes.

The identification of SDPs among protein families can help us understand functional traits that allow diversification among them. This was important to identify key residues to explain differences in substrate selectivity and modes of actions in fungi polygalacturonases GH28 family [20]. As such, we focused our analysis on the identification of SDPs, under the hypothesis that the identified positions may help us to understand how different families acquired novel functions by, for example, a set a residues that may help to accommodate new acceptors and donors in the binding pocket. However, after running general comparative analysis (**Figure 6** and Supplementary Figure 5), or subfamily dedicated analysis with the HCT/HQT group (**Figure 7**, Supplementary Figure S8 and Supplementary Dataset S6), only a handful of the SDPs were located in the proximity (<6 Å) from the substrates. Also, very low amount of SDPs implicate important different physicochemical properties of SDPs that can explain functional diversification by themselves. Because of the deep understanding on both biochemical and structural details on HCT/HQT enzymes, we chose the CcHCT (from G4-1) with p-coumaroyl-CoA and shikimate as reference for the study. Of course, enzymes from different families, showing preference for different substrates, may therefore have binding sites with different architectures allowing the accommodation of the highly dissimilar metabolites that BAHD enzymes can process. Therefore, in a comparative analysis with structures from references in different BAHD groups, we show that most SDPs that are close to substrates in CcHCT are also proximal in other references (Supplementary Figure S4). However, three loops in AtHCT (Q9FI78), corresponding to positions 30-40 (including turn T3-4), 351-372 (T27-28) and 391-401 (T31-32) in CcHCT, are reorganized in the apo-enzyme (compared to the holo-enzyme), generating a shrinkage of the binding pocket [47]. This volume reduction in the active site area was observed also for other HCTs. In addition, the presence of off-center binding sites in HCTs positively affects reaction rate [28]. These sites implicate an evolutionary advantage towards promiscuity and eventually functional diversification. In addition, a recent study identified residues W36 and C320 in Dm3MaT3 as key for the enzyme’s activity [24]. This enzyme corresponds to A4PHY4 from *Chrysanthemum morifolium* in our dataset, which belongs to G2, and we indeed identified both these positions as SDPs (corresponding to SDPs 31 and 302) in different analyses. Both SDPs were identified in the general comparison (**Figure 6**) and are conserved in at least 85% of the sequences in G2’s BAHDome. With this in mind, our method can predict residues that may have distinct roles in different families, although the full mechanism of these features is yet to be established. Potentially, a network of co-evolving SDPs located in different regions of the protein may be responsible for allowing the enzymes to evolve promiscuity (e.g., substrate permissiveness) and consequently, functional diversification. This network of SDPs could have an implication on the protein dynamics, rather than strictly substrate accommodation in the active site, allowing substrate entry and interaction with the binding site(s). Indeed, we were able to identify SDPs subnetworks with residues showing large MI values, even though they are not located proximal in the protein 3D structure.

## FUTURE PERSPECTIVES

Following an approach combining sensitive and specific data mining and clustering, we have generated the BAHDome from 218 land plants. This BAHDome has been clustered in 16 groups. However, we foresee that additional, successive clustering is required to identify actual functional families. In this sense, we show that a not whole group G4, but actually a subgroup G4-1, correspond to a potential functional (sub)family with HCT/HQT activity. In this sense, inception analysis of this subgroup may allow identification of two sub-subfamilies HCT and HQT. Following a similar approach, all identified groups 100P&R-SD can be subjected to such analyses and study in depth evolutionary mechanisms behind the evolution of different BAHD families and subfamilies. In this sense, groups with large number of duplicate homologues per species (e.g., G1) can be under concerted evolution. In addition, we identify several SDPs comparing the different families or a functional subfamily to all other families. In either case, only a handful of SDPs are linked to the actual substrate binding pocket. Therefore, studies of short term protein dynamics can allow us to understand how these residues may be implicated in substrates recognition and accommodation. In addition, a systematic study of the biochemical properties of the enzymes and mutants on these sites can shed light on their implication over activity. Together, these results can be used to successfully classify the complex land plant BAHDome and eventually predict currently unknown activities.

## MATERIALS AND METHODS

### Sequence Mining

#### Training Dataset

A hmmer profile describing the BAHD superfamily was obtained from Pfam[48] (PF02458, Transferase family [49]). This profile was used to run hmmsearch, from HMMER [50], against 17 complete land plant proteomes (Supplementary Dataset S1). In addition, we performed the search against the SwissProt database, limited to Viridiplantae (taxid: 33090). In all cases, we accepted all sequences with a score above the inclusion threshold. The resulting SwissProt dataset was then completed with BAHD sequences described elsewhere [21]. Each dataset was then aligned using MAFFT v7, settings for G-INS-i [51]. To each dataset, we incorporated a reference sequence with 3D structure resolved, HCT from *Coffea canephora* (PDB id 4G0B, [52]) using MAFFT-add. To avoid the high variability in the N- and C-end regions, the multiple sequence alignment was trimmed in these regions according to the first and last residues corresponding to secondary structure in the reference sequence. Subsequently, the processed datasets were subjected to sequence scrutiny using Seqrutinator [53]. Next, in order to avoid overzealous scrutiny, we applied a recovery strategy. For this, the sequences removed by Seqrutinator modules NHHR, GIR y PR (i.e., avoiding modules SSR and CGSR which typically target partial sequences) were combined and clustered by identity (40%) using CD-HIT [54]. The clusters with >10 sequences and with a rather wide taxonomic representation were thus recovered. Next, duplicated sequences, i.e. 100% identity and same length, were removed. These sequences typically derive from different predicted gene models from the same locus. Finally, all sequences from each species were aligned and, upon visual inspection, sequences lacking the catalytic motif HxxxD were removed. This original training sequence set was used to reconstruct a maximum likelihood phylogeny (see below) and subsequently used to generate a clustering (see below). Finally, in order to ensure a complete training dataset, sequences removed by Seqrutinator by modules NHHR, GIR and PR and not recovered were classified into the clustered training dataset, using HMMERCTTER. This results in the final training dataset.

#### Target Dataset

The sequence mining for the target dataset was performed in a similar manner as described for the training. First, we retrieved additional 191 complete land plant proteomes, based on the available sequenced genomes from species described in [55]. The dataset includes proteomes from Gymnosperms (order Pinales=1 species), Eudicots (Proteales=2, Ranunculales=2, Caryophyllales=4, Ericales=3, Gentianales=2, Lamiales=6, Solanales=18, Apiales=2, Asterales=3, Saxifragales=4, Sapindales=8, Malvales=5, Myrtales=2, Brassicales=24, Cucurbitales=8, Fabales=21, Fagales=13, Malpighiales=10, Oxalidales=1, Rosales=22), Monocots (Alismatales=1, Dioscoreales=1, Asparagales=1, Poales=23, Zingiberales=1) and other Angiosperms (Nymphaeales=1, Laurales=1, Magnoliales=1) (see also Supplementary Dataset S5 for more details). For each dataset, we removed i) sequences with non-IUPAC characters and ii) duplicated sequences (100% identity). Next, we performed a hmmsearch on each proteome using the Transferase hmmer profile PF02458. As previously described, each dataset was aligned and N- and C-end trimmed based on the secondary structure of 4G0B. The tool Seqrutinator was applied to each resulting dataset to remove non functional homologues. Finally, all BAHD homologue by each species were combined in a single file, configuring the complete target dataset.

The taxonomical identity of the species used on this study was based on the APG IV angiosperm classification [56] and the NCBI taxonomy database PMC7408187.

The sequences and tree of the complete BAHDome dataset are available as Supplementary Files S2 and S3 (respectively) and Supplementary Dataset S7.

### Phylogenetic reconstruction and statistical support

For small datasets (< 2000 sequences, e.g. the training dataset), the sequences were aligned with MAFFT G-INS-i. Then the reliable columns of the MSA were selected by successively apply trimAl v1.2 (setting -gappyout) [57] and BMGE v1.0 (entropy setting 0.8) [58]. The pruned MSA was used to reconstruct the maximum likelihood phylogeny using PhyML v3.3.3 [59]. We perform the statistical support for the original training phylogeny by generating 1000 botstraps using SeaView v4.0 [60]. This dataset was used to generate the corresponding trees with FastTree v2.1.11 [61]. The branch support was analyzed using transfer bootstrap expectation (TBE) based on Booster [62] calculation and the PhyML tree as reference. For large datasets (i.e., the dataset after target classification), the sequences were aligned using FAMSA2 and the maximum likelihood phylogeny was reconstructed with FastTree.

In all cases, phylogenetic reconstructions were performed using the same parameters (substitution model WAG, gamma shape parameter = 4). Dendroscope v3.7.2 [63] and iTol [64] were used to represent and annotate phylogenetic trees.

### HMMERCTTER clustering and classification

A phylogeny and the unaligned sequences used to generate it were used as inputs for the *clustering* module of HMMERCTTER [65], consistently allowing the identification of clusters with 100% precision and recall self detection (100P&R-SD). In addition, HMMERCTTER *classification* module was used to incorporate sequences from a target dataset into a training set (consisting of sequences in groups 100P&R-SD). As such, the sequences are classified with the condition that the resulting dataset maintain the condition of groups 100P&R-SD.

### 3D structure modelling and visualization

When needed, tridimensional protein structure models (PpaHCT and MpoHCT) were generated using Colabfold [66], applying AlphaFold2 [13] with number of recycles = 5 and amber relaxation activated. AlphaFold2 models for representative SwissProt entries were obtained from Uniprot database. *Coffea canephora* HCT (CcHCT, Uniprot id A4ZKE4) with acceptor (shikimate) and donor (coumaroyl-CoA) was obtained [28]. Pymol [67] was used to visualize and style structures. MSA depicted in Supplementary Fig S1 was generated using NCBI MSA Viewer [68].

### BAHD donor and acceptor metabolites multidimensional scaling and clustering

We investigated the literature to generate a list of the metabolites (both donors and acceptors) that have been described for the SwissProt entries identified in out datamining. We accepted only those metabolites that have been confirmed at the biochemical level, with at least one kinetic parameter measured (e.g., KM or Vmax). When kinetic properties for several metabolites are measured for a given SwissProt entry, we only considered those values ≥20% of the maximal value. Then, we obtained the SMILES formatted files for all metabolites (93 acceptors and 21 donors) from PubChem database [69], except for nine metabolites which were manually annotated using Marvin JS [70] and exported as SMILES. Next, we perform similarity search with ChemMine tools [71] for donors and acceptors (separately), followed by the generation of 3D vectors of the chemical compounds using the multidimensional scaling 3D by ChemMine clustering tool. These vectors condense data on several similarity coefficients, including structural descriptors and Tanimoto coefficient. The resulting 3D vectors were clustered by Kmeans (K=12 and =4 for acceptors and donors, respectively) with Scikit-learn package in Python [72]. Data was plotted in 3D charts using plotly [73].

### Comparison of percentage of identity among sequences

All the sequences in one group were compared to all the sequences in such group (i.e., internal comparison) and to all the sequences in each of the remaining groups. To do so, each sequence pair was aligned by Needleman-Wunsch alghoritm, using global:global ggsearch program from FASTA (v36.3.8h) [74]. The percentage of identity (ID%) was recorded for each pair. The distribution of the ID% was plotted using plotly. The datasets were analyzed with JASP v0.16.3 [75] to calculate mean, quartiles, Skewness and Kurtosis.

### Identification of potential clusters of orthologues in BAHDome

We perform a search of groups of orthologues in the complete BAHDome (training+classified target sequences) using incremental clustering based on bidirectional best BLAST hit (BDBH), implemented in GET_HOMOLOGUES_EST program [76]. In order to ensure best quality datasets, we did not include sequences from orders with very low homologue number comparing neighbor orders (such as Oxalidales from Eudicots; Asparagales, Dioscoreales, Zingiberales from Monocots; Ginkgoales, Cupressales from Gymnosperms; Marchantiales, Funariales from Bryophytes). The sequences were then grouped by Order, resulting in a total of 15715 sequences in 29 Orders. Two different analyses were performed: identification of orthologues in (i) all described Orders (ii) only Eudicot and Monocot Orders. In each case, the analysis was performed enabling Pfam and BDBH searches. In addition, and considering the variability of BAHD homologues (even in the same group), we decreased the minimum percentage of identity to 50. For the resulting BDBH clusters, the sequences ids were retrieved and located in the phylogeny reconstructed with the classified dataset.

### Generation of SDP networks and representation using sequence logos

Sequences from the groups to compare were aligned with MAFFT G-INS-i. The MSA was then used to identify clustering determining positions (CDPs) using SDPfox module *sdplight* [17]. The same MSA was used to determine MI using Mistic [18]. With these inputs, we used a dedicated Python script to generate an SDP network (SN). The basic idea of the SN is that highly interconnected CDPs are less likely to have resulted from drift and, as such, more likely to be SDP. The script takes all CDPs determined by SDPfox with z-score > 3.29, and all MI z-scores > 3.29 according to Mistic. For a given MI threshold (MIT), the Mutual Information Connectivity Score (MICS) of a CDP*i* expresses its total MI to other CDPs (the sum of its z-scores) divided by the number of connections the CDP*i* node has to other CDP-nodes:

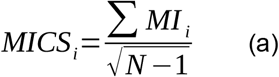

where N = number of CDPs that have at least one connection with z-score large than the MIT.

The script determines a range of MICSs for all CDPs with MITs in between 3 .29 and the highest MI z-score of the CDPs. Note that the MICS of a node as such will increase until the corresponding CDP is excluded by the MIT.

Then, an SN score or Specific Network Score *SNS* is calculated for each MIT with the formula

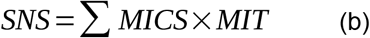

By increasing the MIT, the number of nodes in the network will decrease, and the SNS will typically increase until a maximum is reached, followed by a progressive decrease in the SNS. As a result the SNS is a compound measure that weighs the amount of connections to the strength of the connections. We therefore select the best SN as that presenting the maximal SNS. Once the SN is determined, the script ensembles 1000 random networks with the same node number as the SN, using all nodes (both CDPs or not CDPs) and the same cut-offs. This implicates that by chance, random networks may or may not contain actual CDPs. Then, the SNS for each random network is calculated as well as the z-score for the SN in the distribution of the random generated networks.

For SDP logos, a custom script was used to isolate SDP columns into pseudo SDP MSAs. Sequence logos (both to display bits and calculate residues frequencies per residue) were made using weblogo [77].

## Notes

### Competing Interest Statement

The authors have declared no competing interest.

