## Supplementary Figures S1-S9 for "Comprehensive characterization of the complex BAHD acyltransferase family from 218 land plants species: phylogenomic analysis and identification of specificity determinant positions"

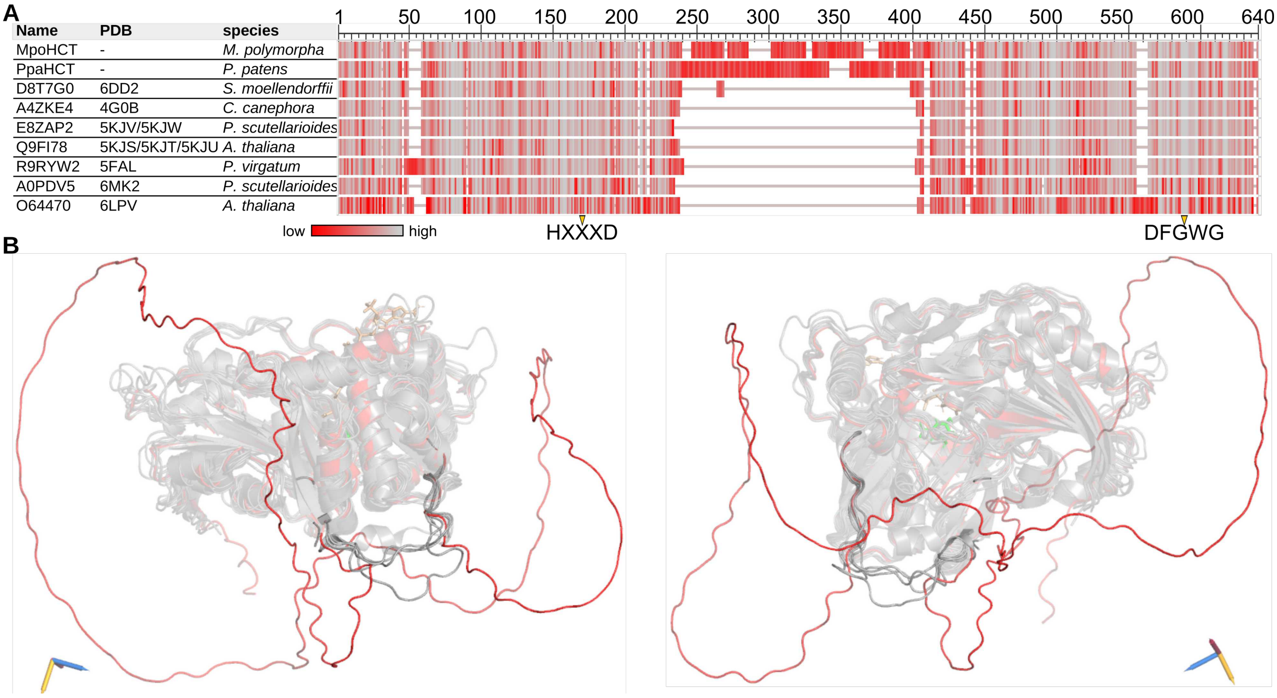


**Supplementary_Figure 1**: *MpoHCT and PpaHCT present a large insert*. (A) Seven sequences from G4 with resolved 3D structures were aligned to PpaCT and MpoHCT. The residues are colored according to the shown scale from low to high Column Quality Score (using MSA viewer @ NCBI). The regions corresponding to classic motifs HxxxD and DFGWG are indicated. (B) The structure of PpaHCT (in red) was structurally aligned to the resolved structures of several HCT/HQT (shown in gray). The region of the loop corresponding to the insert in PpHCT is highlighted in all structures. Two rotational views are shown, as indicated by the coordinate axes. RMSD values (in Å) to CcHCT are: PpHCT 0.664 (187 atoms), 6DD2 0.781 (330 atoms), 5KJV 0.462 (365 atoms), 5KJS 0.506 (362 atoms), 5FAL 1.071 (385 atoms), 6MK2 1.133 (353 atoms), 6LPV 1.151 (309 atoms).

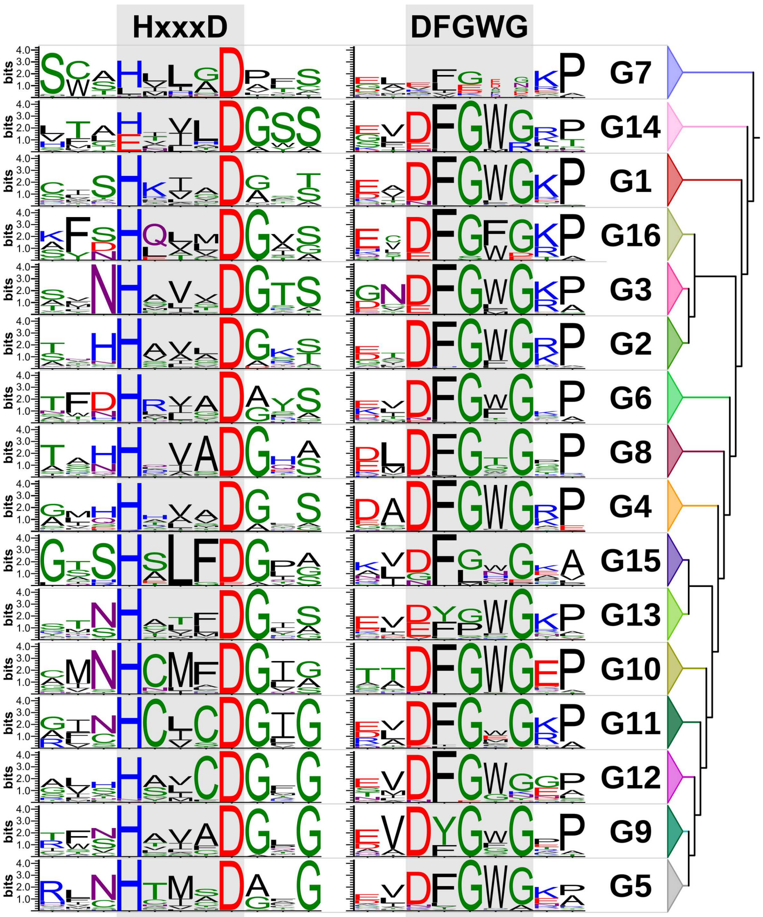


**Supplementary_Figure 2**: *Conservation of BAHD classic motifs in G7 and other training groups*. Sequence logos depict conservation of HxxxD motif (with three additional residues to the N- and C-end) and DFGWG motif (two additional residues) for the sequences in each group. Groups are organized by their phylogenetic relationships.

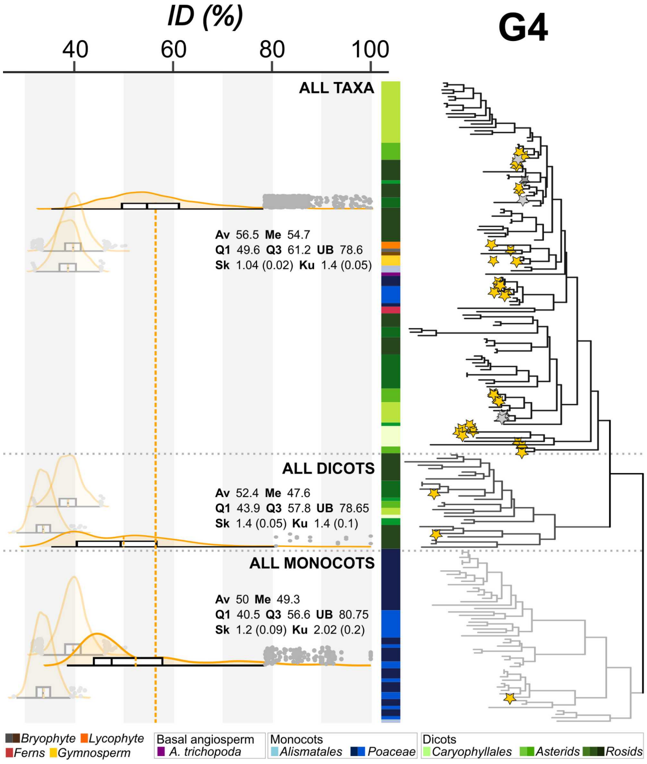


**Supplementary_Figure 3:** *G4 shows internal topology with taxonomical division*. The three subclades in G4 are indicated as ALL TAXA, ALL MONOCOTS and ALL DICOTS. The stars indicate SP entries. The taxa for each leaf is indicated with the colored bar, using the coloring indicated below. To the left, distribution of ID% among all sequences in one subgroup compared to all sequences in each subgroup. Distributions plots are shown as split violins with its corresponding boxplot. Comparisons of sequences in one subgroup with themselves (internal comparison) are shown with maximum opacity. Dashed lines, distribution average (extended line is average of the internal comparison in the ALL TAXA subgroup). Outliers points are also shown. The order of the distribution plots correspond is the same top to bottom: ALL TAXA, ALL MONOCOTS and ALL DICOTS. Statistical values for internal comparisons are shown: Av, Average; Q1, first quartile; Me, Median; Q3, third quartile; UB, upper bound; Sk, Skewness (Standard Error); Ku, Kurtosis (Standard Error).

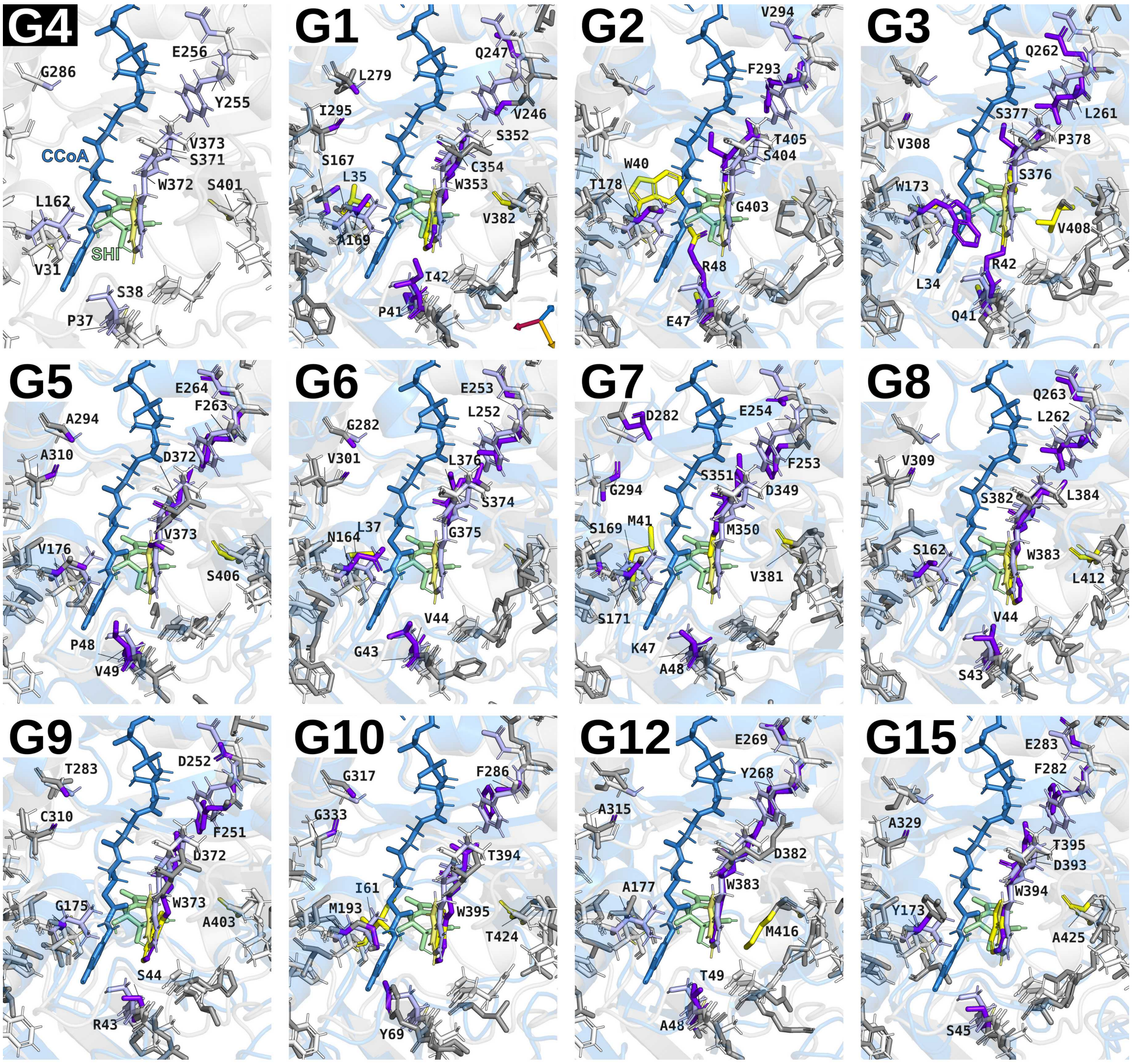


| | Uniprot id | RMSD | SDP | | | | | | | | | | | |
|---|---|---|---|---|---|---|---|---|---|---|---|---|---|---|
| ***G4*** | *A4ZKE4* | - | *31* | *37* | *38* | *160* | *162* | *255* | *256* | *286* | *302* | *371* | *372* | *373* | *401* |
| **G1** | A0A2P1GIW7 | 1.82 | 35 | 41 | 42 | 167 | 169 | 246 | 247 | 279 | 295 | 352 | 353 | 354 | 382 |
| **G2** | Q940Z5 | 2.00 | 40 | 47 | 48 | | 178 | 293 | 294 | | | 403 | 404 | 405 | |
| **G3** | Q9SND9 | 1.38 | 34 | 41 | 42 | | 173 | 261 | 262 | | 308 | 376 | 377 | 378 | 408 |
| **G5** | Q6E593 | 1.62 | | 48 | 49 | | 176 | 263 | 264 | 294 | 310 | 372 | 373 | | 406 |
| **G6** | A1XWY7 | 1.98 | 37 | 43 | 44 | | 164 | 252 | 253 | 282 | 301 | 374 | 375 | 376 | |
| **G7** | Q9LIS1 | 2.90 | 41 | 47 | 48 | 169 | 171 | 253 | 254 | 282 | | 349 | 350 | 351 | 381 |
| **G8** | Q7XPK7 | 1.13 | | 43 | 44 | | 162 | 262 | 263 | | 309 | 382 | 383 | 384 | 412 |
| **G9** | Q5JNT2 | 2.19 | | 43 | 44 | | 175 | 251 | 252 | 283 | 310 | 372 | 373 | | 403 |
| **G10** | Q94CD1 | 1.52 | 61 | | 69 | | 193 | 286 | | 317 | 333 | 394 | 395 | | 424 |
| **G12** | Q8GYW8 | 1.50 | | 48 | 49 | | 177 | 268 | 269 | | 315 | 382 | 383 | | 416 |
| **G15** | Q9SZ58 | 1.37 | | | 45 | | 173 | 282 | 283 | | 329 | 393 | 394 | 395 | 425 |

**Supplementary_Figure 4**: *Structural localization of homologous SDPs in BAHD references from different groups*. Reference SP entry from each group (blue cartoon, gray sticks) is aligned to G4's reference CcHCT A4ZKE4 (gray cartoon, white sticks) with donor coumaroyl-CoA (CCoA) and acceptor shikimate (SHI). For each case, RMSD value is shown. Cartoons are shown with transparency. Sticks for all SDPs are shown, however labeled SDPs from each group reference are those with atoms located at least at 6 Å from the substrates. In addition, SDPs with atoms close to the donor are colored purple, whereas those close to the acceptor are colored yellow (for CcHCT, colors are light purple and light yellow, respectively). In the table, each column correspond to CcHCT homologous SDPs in each reference (residue numbering for each reference structure). Cells are colored according to their proximity to either donor (purple), acceptor (yellow) or mixed (both).

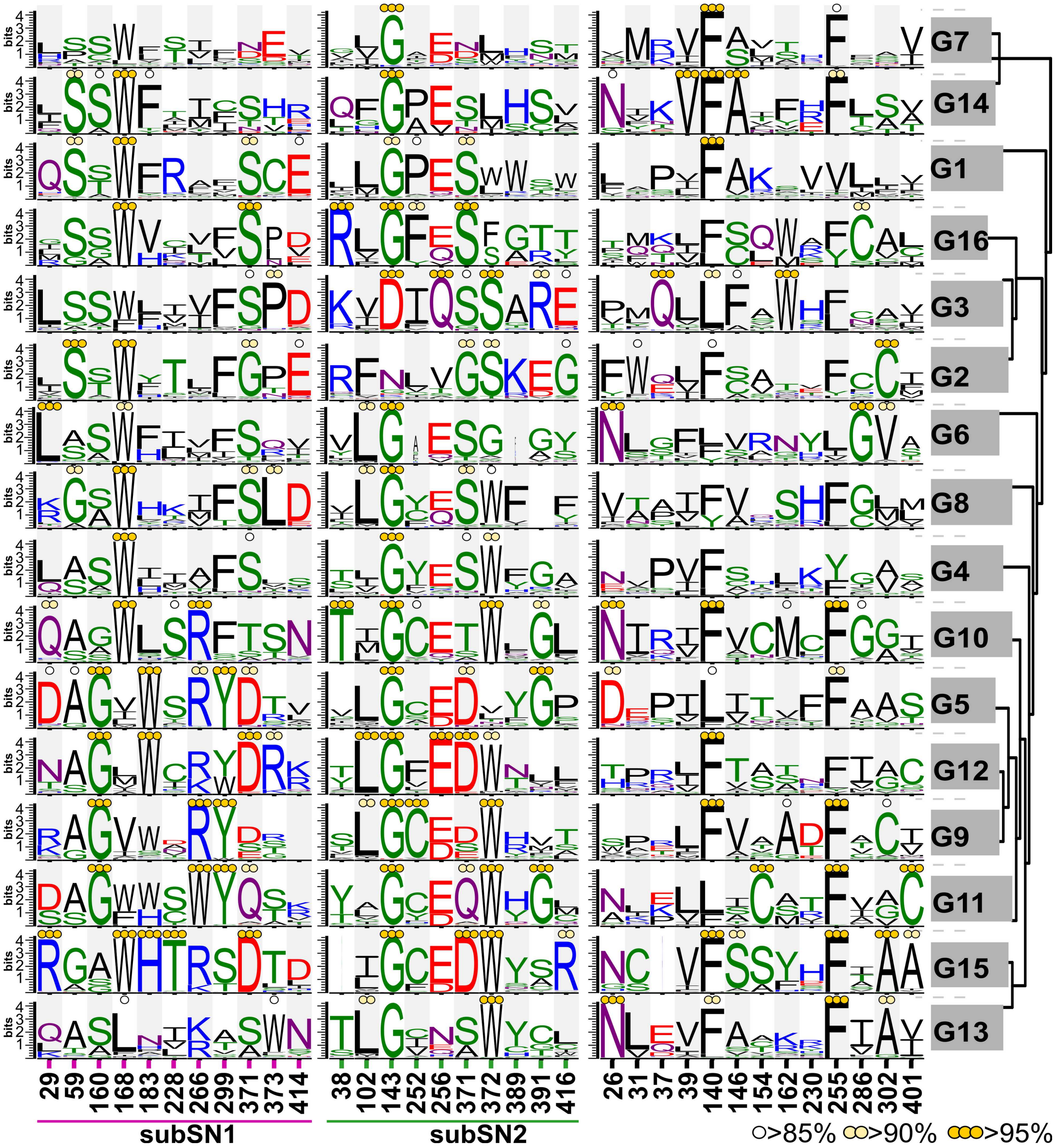


**Supplementary_Figure 5:** *Weblogos of SDPs identified in the families for the complete BAHDome*. The weblogos for the 33 SDPs identified with the training dataset are generated using the complete BAHDome (i.e., the families including the the sequences following target classification). In rows, BAHD families are indicated, arranged according to their phylogenetic relationships. SDPs are numbered according to a reference sequence CcHCT. Note that SDPs are arranged based on their occurrence or not in a subSN. Y-axis show bits. Colored dots indicate SDPs for which either >85%, >90% or >95% of the sequences in the family show a conserved residue.

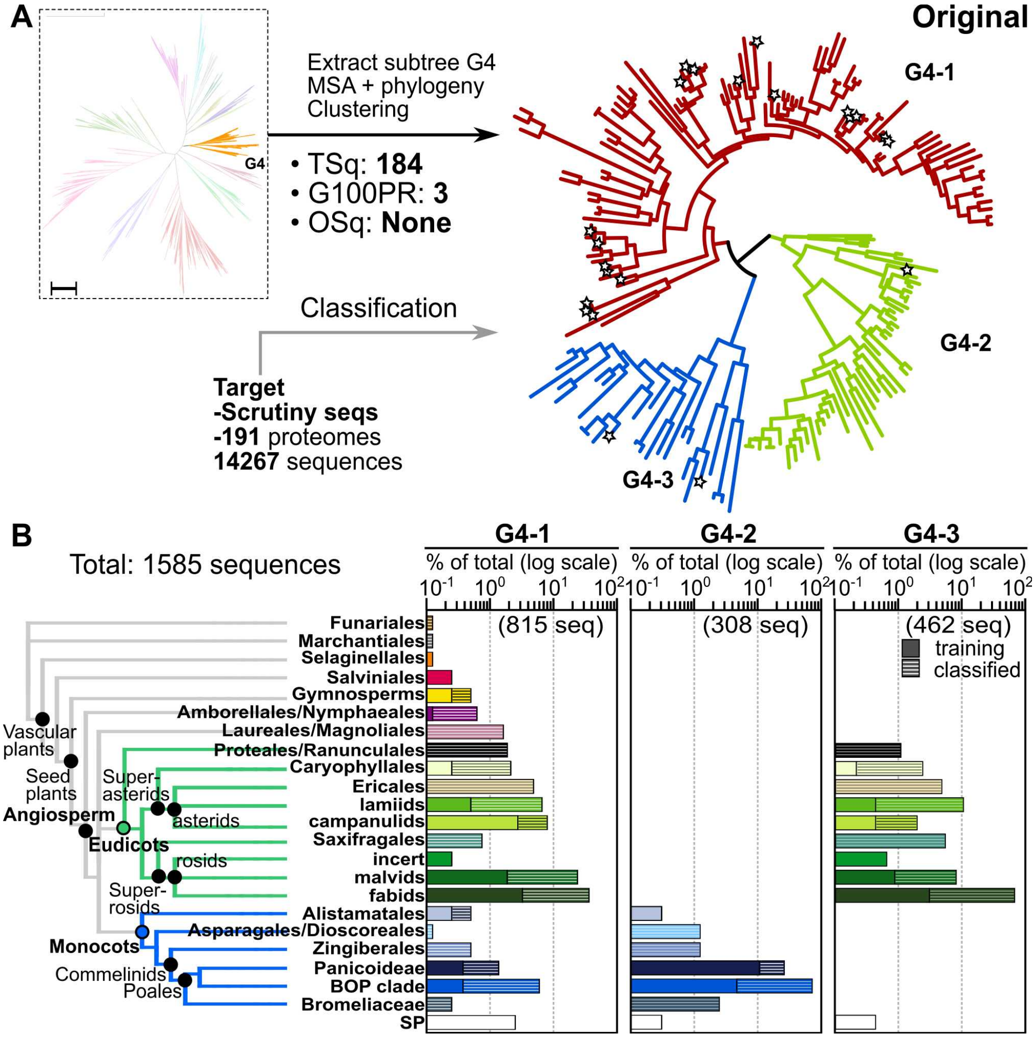


**Supplementary_Figure 6**: *Dedicated clustering and classification analysis for G4 shows three subgroups*. (A) G4 sequences from the original set are extracted and used to reconstruct a dedicated phylogeny followed by HMMERCTTER clustering, resulting in three groups 100P&R-SD. Stars indicate SP entries. TSq: total sequences; OSq: Orphan (not clustered) sequences. (B) Clustered G4 was used to classify the target dataset. The target includes the sequences not recovered and not partial removed by Seqrutinator, plus all BAHD homologues retrieved from 191 land plant proteomes. The relative amount of sequences in each subgroup after classification is shown (in log scale); sequences from original and target sets are indicated in plain or striped bars, respectively. Total number of sequences is shown in brackets for each subgroup.

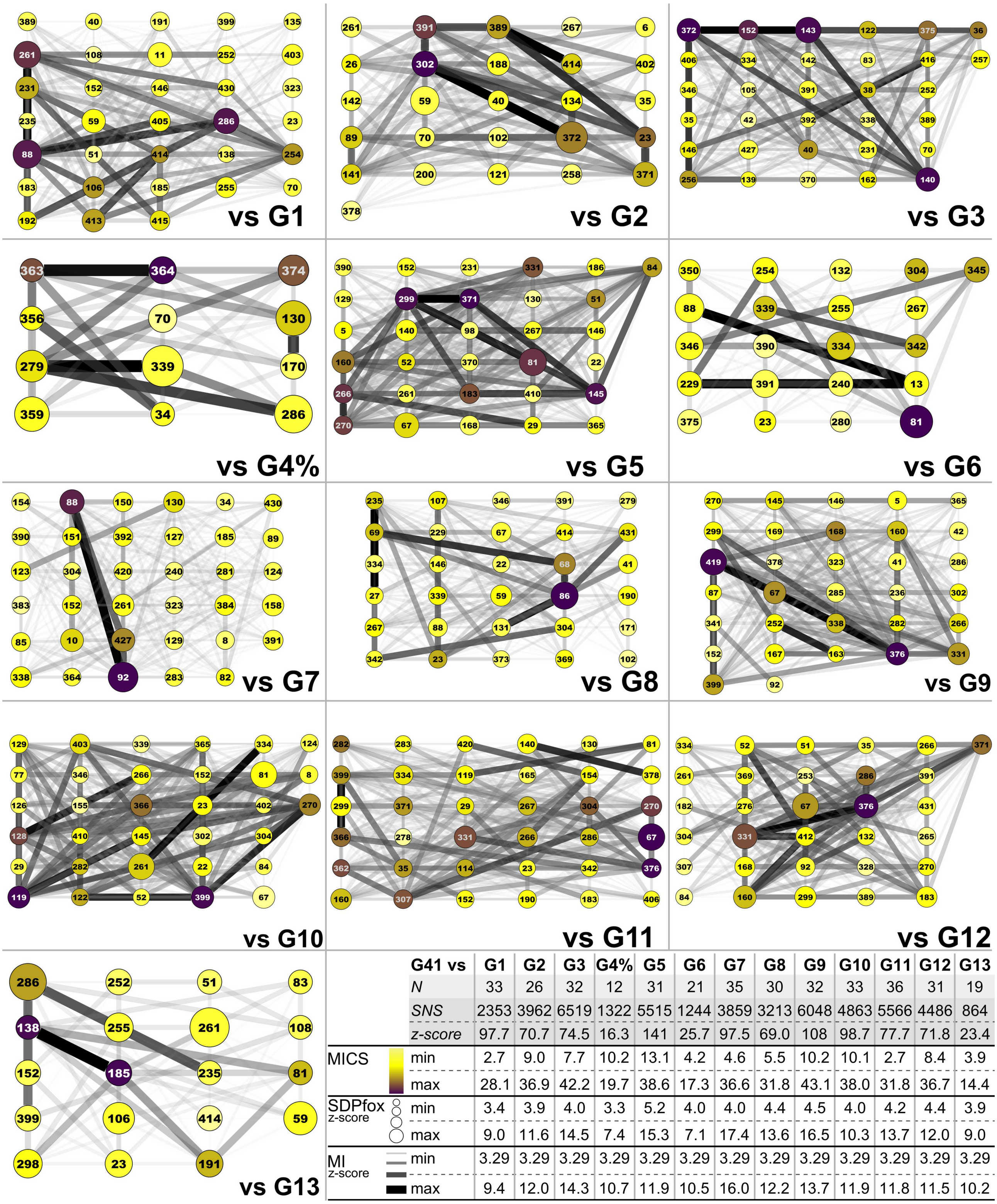


| | G41 vs | G1 | G2 | G3 | G4% | G5 | G6 | G7 | G8 | G9 | G10 | G11 | G12 | G13 |
|---|---|---|---|---|---|---|---|---|---|---|---|---|---|---|
| | *N* | 33 | 26 | 32 | 12 | 31 | 21 | 35 | 30 | 32 | 33 | 36 | 31 | 19 |
| | *SNS* | 2353 | 3962 | 6519 | 1322 | 5515 | 1244 | 3859 | 3213 | 6048 | 4863 | 5566 | 4486 | 864 |
| | *z-score* | 97.7 | 70.7 | 74.5 | 16.3 | 141 | 25.7 | 97.5 | 69.0 | 108 | 98.7 | 77.7 | 71.8 | 23.4 |
| MICS | min | 2.7 | 9.0 | 7.7 | 10.2 | 13.1 | 4.2 | 4.6 | 5.5 | 10.2 | 10.1 | 2.7 | 8.4 | 3.9 |
| | max | 28.1 | 36.9 | 42.2 | 19.7 | 38.6 | 17.3 | 36.6 | 31.8 | 43.1 | 38.0 | 31.8 | 36.7 | 14.4 |
| SDPfox z-score | min | 3.4 | 3.9 | 4.0 | 3.3 | 5.2 | 4.0 | 4.0 | 4.4 | 4.5 | 4.0 | 4.2 | 4.4 | 3.9 |
| | max | 9.0 | 11.6 | 14.5 | 7.4 | 15.3 | 7.1 | 17.4 | 13.6 | 16.5 | 10.3 | 13.7 | 12.0 | 9.0 |
| MI z-score | min | 3.29 | 3.29 | 3.29 | 3.29 | 3.29 | 3.29 | 3.29 | 3.29 | 3.29 | 3.29 | 3.29 | 3.29 | 3.29 |
| | max | 9.4 | 12.0 | 14.3 | 10.7 | 11.9 | 10.5 | 16.0 | 12.2 | 13.7 | 11.9 | 11.8 | 11.5 | 10.2 |

**Supplementary_Figure 7**: *SDP retworks (SNs) resulting from comparison of training group 4-1 vs each of the remaining groups*. The networks represented by SDP (nodes) connected by MI values (edges). G4% indicates the comparison vs the remainder of G4 (G4-2/G4-3). The table indicates on top the number of nodes (N), SDP network score (SNS) and the z-score of the SN compared to the score of 1000 randomly generated networks. For each node, we indicate the MICS value by the color according to the scale and the z-value determined by SDPfox with the sphere diameter. For edges, MI z-score value is indicated by shade and width. For MICS, SDPfox and MI z-scores, minimum and maximum values for each case is shown. Node label correspond to position on the reference sequence CcHCT.

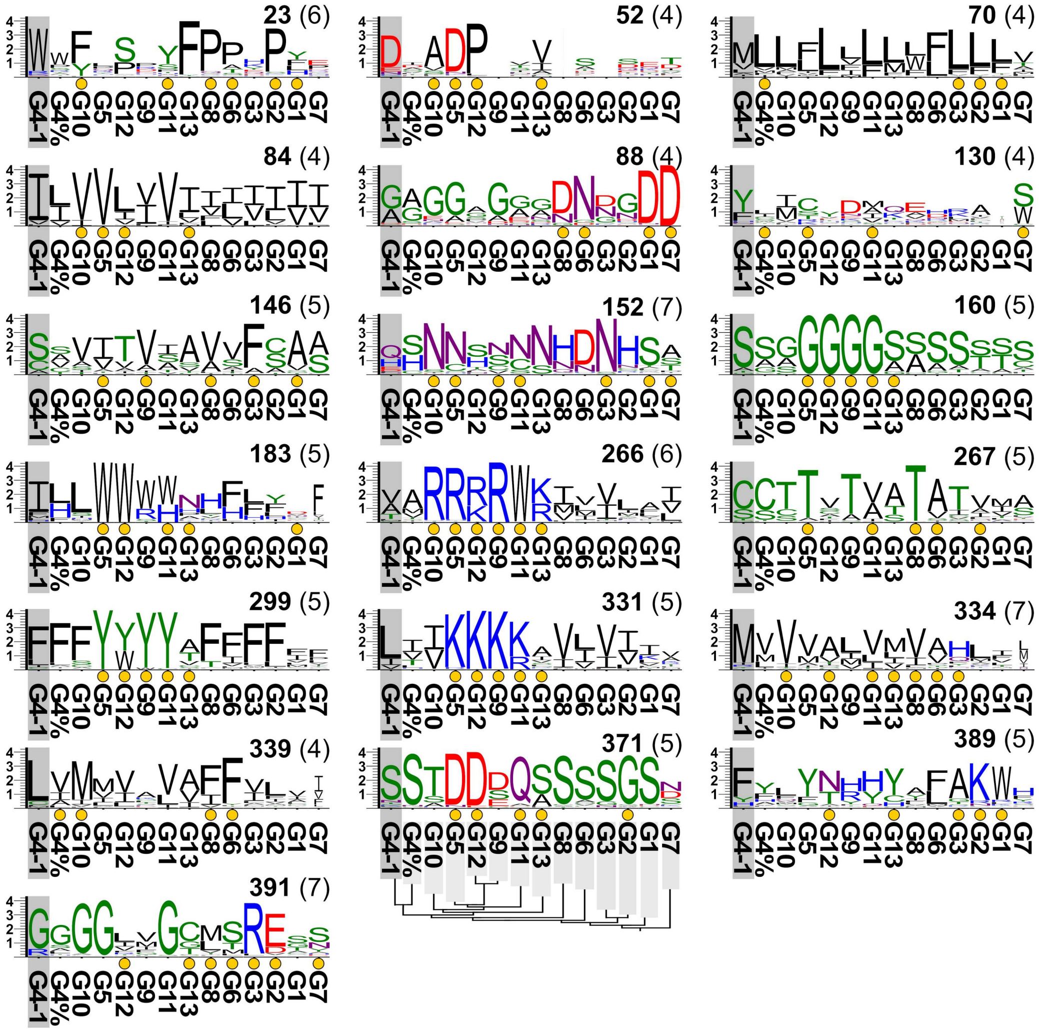


**Supplementary_Figure 8**: *Sequence logos of SDPs identified in at least four comparisons and conserved in <90% of G4-1's sequences after target classification*. SDPs are indicated in top right (in brackets the number of comparisons in which the SDP was found). Yellow dots indicate comparisons in which the position was actually identified as an SDP. The phylogenetic relationship among groups is shown. Logos x-axes values indicate bits. G4% indicates the remainder of G4, i.e. sequences in G4 not belonging to G4-1.

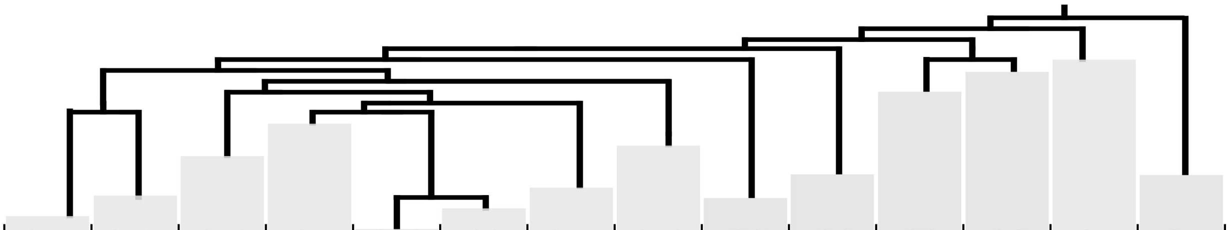

| SDP | G41 | G4% | G10 | G5 | G12 | G9 | G11 | G13 | G8 | G6 | G3 | G2 | G1 | G7 |
|---|---|---|---|---|---|---|---|---|---|---|---|---|---|---|
| Seqs | 801 | 794 | 576 | 1540 | 479 | 628 | 642 | 307 | 654 | 825 | 2310 | 1922 | 3327 | 631 |
| 23 | 86% (W) | 48% (W) | 70% (F) | 22% (L) | 63% (S) | 27% (P) | 58% (Y) | 95% (F) | 87% (P) | 63% (P) | 26% (H) | 88% (P) | 32% (Y) | 30% (E) |
| 35 | 92% (H) | 81% (H) | 60% (I) | 45% (Q) | 26% (T) | 32% (L) | 36% (S) | 34% (D) | 33% (H) | 50% (D) | 67% (I) | 41% (P) | 68% (Y) | 50% (Y) |
| 51 | 91% (F) | 43% (F) | 30% (D) | 37% (K) | 67% (D) | | 27% (V) | 22% (I) | | 21% (G) | | 16% (L) | 17% (S) | 43% (L) |
| 52 | 75% (D) | 18% (I) | 75% (A) | 88% (D) | 93% (P) | | 22% (V) | 62% (V) | 1% (V) | 40% (S) | | 29% (S) | 28% (E) | 35% (T) |
| 67 | 95% (F) | 76% (F) | 93% (Y) | 94% (Y) | 92% (Y) | 93% (Y) | 99% (Y) | 99% (Y) | 80% (Y) | 56% (Y) | 89% (F) | 66% (F) | 65% (F) | 58% (Y) |
| 70 | 60% (M) | 77% (L) | 76% (L) | 50% (F) | 85% (L) | 40% (L) | 74% (L) | 65% (L) | 40% (L) | 64% (F) | 83% (L) | 77% (L) | 60% (L) | 43% (V) |
| 81 | 94% (R) | 96% (R) | 90% (K) | 94% (K) | 55% (R) | 30% (E) | 63% (K) | 97% (R) | 48% (R) | 78% (E) | 37% (S) | 57% (K) | | 49% (R) |
| 84 | 89% (I) | 50% (L) | 89% (V) | 97% (V) | 64% (L) | 51% (V) | 91% (V) | 65% (I) | 47% (I) | 41% (I) | 49% (I) | 60% (I) | 59% (I) | 55% (I) |
| 88 | 58% (G) | 51% (A) | 75% (G) | 76% (G) | 32% (A) | 76% (G) | 42% (G) | 40% (G) | 72% (D) | 88% (N) | 35% (D) | 59% (G) | 95% (D) | 99% (D) |
| 92 | 94% (L) | 38% (L) | 69% (V) | 53% (L) | 88% (P) | 57% (W) | 52% (L) | 49% (G) | 59% (R) | 48% (D) | 36% (E) | 55% (S) | 24% (E) | 100% (R) |
| 130 | 62% (Y) | 25% (L) | 40% (I) | 44% (C) | 35% (Y) | 51% (D) | 26% (M) | 33% (Q) | 41% (E) | 25% (H) | 24% (R) | 48% (A) | 4% (T) | 54% (S) |
| 146 | 66% (S) | 27% (S) | 64% (V) | 58% (I) | 66% (T) | 78% (V) | 33% (I) | 69% (A) | 69% (V) | 53% (V) | 87% (F) | 41% (C) | 78% (A) | 54% (A) |
| 152 | 47% (Q) | 51% (S) | 90% (N) | 75% (N) | 41% (S) | 39% (N) | 61% (N) | 81% (N) | 65% (H) | 78% (D) | 95% (N) | 75% (H) | 75% (S) | 44% (A) |
| 160 | 85% (S) | 60% (S) | 56% (G) | 99% (G) | 100% (G) | 99% (G) | 99% (G) | 69% (S) | 52% (S) | 73% (S) | 79% (S) | 45% (S) | 45% (S) | 58% (S) |
| 168 | 99% (W) | 100% (W) | 100% (W) | 45% (V) | 35% (L) | 80% (V) | 65% (W) | 86% (L) | 99% (W) | 95% (W) | 71% (W) | 99% (W) | 98% (W) | 84% (W) |
| 183 | 75% (I) | 51% (L) | 75% (L) | 99% (W) | 100% (W) | 68% (W) | 62% (W) | 49% (N) | 53% (H) | 62% (F) | 41% (L) | 32% (Y) | 22% (D) | 37% (F) |
| 261 | 97% (H) | 80% (H) | 80% (F) | 53% (C) | 50% (Y) | 37% (K) | 97% (H) | 45% (H) | 97% (H) | 46% (F) | 48% (H) | 61% (Y) | 54% (L) | 55% (L) |
| 266 | 42% (V) | 57% (A) | 96% (R) | 94% (R) | 63% (R) | 99% (R) | 98% (W) | 56% (K) | 51% (I) | 34% (V) | 54% (V) | 46% (L) | 35% (A) | 43% (I) |
| 267 | 68% (C) | 73% (C) | 67% (T) | 91% (T) | 41% (V) | 82% (T) | 51% (V) | 58% (A) | 97% (T) | 75% (A) | 65% (T) | 48% (V) | 32% (M) | 46% (A) |
| 270 | 92% (R) | 99% (R) | 96% (L) | 89% (L) | 55% (L) | 37% (I) | 93% (L) | 21% (V) | 99% (R) | 26% (A) | 80% (R) | 33% (R) | 30% (R) | 42% (R) |
| 286 | 94% (G) | 30% (I) | 89% (G) | 61% (A) | 70% (I) | 29% (A) | 45% (V) | 69% (I) | 70% (G) | 96% (G) | 38% (C) | 60% (C) | 67% (L) | 20% (F) |
| 299 | 76% (F) | 85% (F) | 79% (F) | 99% (Y) | 59% (Y) | 96% (Y) | 97% (Y) | 43% (A) | 81% (F) | 64% (F) | 84% (F) | 82% (F) | 30% (F) | 35% (F) |
| 304 | 95% (F) | 24% (L) | 85% (V) | 57% (A) | 64% (V) | 37% (Y) | 68% (V) | 59% (L) | 97% (L) | 72% (S) | 46% (Q) | 21% (G) | 22% (W) | 36% (V) |
| 331 | 74% (L) | 41% (I) | 54% (I) | 95% (K) | 99% (K) | 98% (K) | 73% (K) | 43% (A) | 77% (V) | 43% (L) | 71% (V) | 54% (I) | 41% (I) | 28% (V) |
| 334 | 85% (M) | 53% (V) | 87% (V) | 62% (V) | 62% (A) | 47% (L) | 62% (V) | 44% (M) | 71% (V) | 53% (A) | 56% (H) | 55% (L) | 27% (I) | 13% (L) |
| 339 | 85% (L) | 57% (V) | 82% (M) | 45% (M) | 58% (V) | 35% (A) | 70% (V) | 53% (A) | 68% (F) | 90% (F) | 46% (V) | 52% (L) | 30% (V) | 20% (I) |
| 346 | 90% (L) | 58% (L) | 82% (F) | 68% (M) | 39% (Q) | 1% (L) | 52% (V) | 58% (G) | 53% (A) | 71% (V) | 91% (W) | 19% (L) | 26% (L) | 26% (L) |
| 371 | 81% (S) | 94% (S) | 71% (T) | 94% (D) | 95% (D) | 49% (D) | 91% (Q) | 67% (S) | 92% (S) | 80% (S) | 85% (S) | 92% (G) | 91% (S) | 36% (N) |
| 376 | 91% (P) | 84% (P) | 57% (S) | 76% (G) | 89% (G) | 99% (G) | 72% (G) | 81% (G) | 53% (P) | 80% (P) | 30% (N) | 39% (G) | 62% (P) | 32% (D) |
| 389 | 71% (F) | 42% (Y) | 38% (L) | 71% (Y) | 56% (N) | 44% (H) | 55% (H) | 67% (Y) | 35% (A) | 54% (F) | 75% (A) | 82% (K) | 79% (W) | 38% (H) |
| 391 | 83% (G) | 62% (G) | 94% (G) | 97% (G) | 37% (L) | 38% (V) | 96% (G) | 55% (C) | 49% (M) | 60% (S) | 93% (R) | 76% (E) | 36% (S) | 50% (S) |
| 399 | 97% (G) | 88% (G) | 89% (E) | 38% (G) | 37% (E) | 52% (P) | 30% (R) | 54% (D) | 99% (G) | 78% (G) | 97% (G) | 53% (G) | 70% (N) | 98% (G) |

Colour scale: 0 10 20 30 40 50 60 70 80 90 100

% of group sequences with a conserved residue (maximal)

**Supplementary_Figure 9**: *SDPs identified are conserved at different ratios in the different BAHD families*. The proportion of sequences presenting an specific residue corresponding to the identified SDP position was calculated. The residue represented in the maximal proportion of sequences in a family is shown in brackets. The cell background is colored according to the scale. The number of sequences in each family is also shown. G4% indicates the remainder of G4, i.e. sequences in G4 not belonging to G4-1.
